# The necrotrophic effector ToxA from *Parastagonospora nodorum* interacts with wheat NHL proteins to facilitate *Tsn1*-mediated necrosis

**DOI:** 10.1101/2021.09.22.461440

**Authors:** Bayantes Dagvadorj, Megan A. Outram, Simon J. Williams, Peter S. Solomon

**Affiliations:** Research School of Biology, The Australian National University, Canberra, ACT 2601, Australia

**Author notes:** **Corresponding author: Peter S. Solomon**.

**Keywords:** necrotrophic effectors, ToxA, NHL proteins, NDR1, *Parastagonospora nodorum*, wheat

## Abstract

The plant pathogen *Parastagonospora nodorum* secretes necrotrophic effectors to promote disease. These effectors induce cell death on wheat cultivars carrying dominant susceptibility genes in an inverse gene-for-gene manner. However, the molecular mechanisms underpinning these interactions and resulting cell death remain unclear. Here, we used a yeast-two-hybrid library approach to identify wheat proteins that interact with the necrotrophic effector ToxA. Using this strategy, we identified an interaction between ToxA and a wheat transmembrane NDR/HIN1-like protein (TaNHL10) and confirmed the interaction using *in-planta* co-immunoprecipitation and confocal microscopy co-localization analysis. We showed that the C-terminus of TaNHL10 is extracellular whilst the N-terminus was localized in the cytoplasm. Further analyses using yeast-two-hybrid and confocal microscopy co-localization showed that ToxA interacts with the C-terminal LEA2 extracellular domain of TaNHL10. Random mutagenesis was then used to identify a ToxA mutant, ToxA^N109D^, which was unable to interact with TaNHL10 in yeast-two-hybrid assays. Subsequent heterologous expression and purification of ToxA^N109D^ in *Nicotiania benthamiana* revealed that the mutated protein was unable to induce necrosis on *Tsn1*-dominant wheat cultivars confirming that the interaction of ToxA with TaNHL10 is required to induce cell death. Collectively, these data advance our understanding on how ToxA induces cell death during infection and further highlights the importance of host cell surface interactions in necrotrophic pathosystems.

## Introduction

Plant fungal pathogens are causal to severe yield losses in cereal production. These pathogens secrete proteinaceous molecules called effectors, which manipulate and facilitate the colonisation the host plant (Dodds and Rathjen, 2010). Effector molecules have been the subject of intense study over the past decade as their requirement and importance for mediating plant diseases of global importance has been recognised. It is acknowledged that an increased understanding of their function during disease can contribute to the development of novel strategies to manage plant diseases.

The ToxA effector is a small, secreted protein that induces necrosis (cell death) in susceptible wheat cultivars. It is closely associated with the enhanced pathogenicity and virulence of *Pyrenophora triciti-repentis, Bipolaris sorokiniana*, and *Parastagonospora nodorum* (Oliver *et al*., 2012; McDonald and Solomon, 2018; McDonald *et al*., 2018). These pathogens are classic necrotrophic fungi that thrive on dead or dying tissue to complete their infection cycle and it has been demonstrated that effector-induced host cell death is central to the infection mechanisms of these pathogens (McDonald and Solomon, 2018). ToxA only induces cell death in wheat cultivars harbouring the dominant susceptiblility host gene, *Tsn1* (Faris *et al*., 2010). *Tsn1* encodes a protein with nucleotide-binding, leucine-rich repeat, and N-terminal serine/threonine protein kinase domains thus showing similarity to classically described resistance genes that act against biotrophic pathogens. However, the function or mechanism of *Tsn1* is unknown, and there is no evidence that ToxA and Tsn1 directly interact (Faris *et al*., 2010).

There are multiple reports published on deciphering the mechanism of ToxA-induced necrosis through protein interaction studies (Manning *et al*., 2007; Lu *et al*., 2014; Tai *et al*., 2007). Manning et al. (2007) showed that ToxA associates with the chloroplast protein ToxABP1, a homologue to the Thylakoid formation 1 (Thf1) protein from *Arabidopsis thaliana*. It was suggested that the ToxA-ToxABP1 interaction is likely to cause oxidative stress by interrupting the chloroplast physiological function leading to cell death (Ciuffetti *et al*., 2010). Another study used a yeast 2-hybrid approach to identify that ToxA interacts with chloroplast-localised plastocyanin (Tai *et al*., 2007). In this study, the silencing of plastocyanin in wheat showed localized cell-death similar to ToxA-induced necrosis, suggesting that the ToxA-plastocyanin interaction may have a role in facilitating cell death (Tai *et al*., 2007). In a more recent study, ToxA was shown to interact with the pathogenesis-related protein PR1-5 by Y2H screen (Lu *et al*., 2014). Site-directed mutagenesis studies identified essential residues on both ToxA and PR-1-5, which were necessary for interaction and cell death (Lu *et al*., 2014).

Previous mutagenesis studies and a peptide (RGDV) competition assays in ToxA have identified that an arginyl-glycyl-aspartic (RGD, 140-142) tripeptide motif in a solvent-exposed loop is required for *Tsn1*-associated necrosis and ToxA internalization into host cells (Manning *et al*., 2008; Meinhardt *et al*., 2002). Interestingly, in animals, RGD motifs are known to be recognized by animal integral membrane receptors, called integrins, to facilitate endocytosis (Gee *et al*., 2008; Ruoslahti and Pierschbacher, 1986). Thus, it was hypothesized that ToxA binds to a surface-exposed receptor through its RGD motif leading to its internalization. However, to date, no such receptor has been reported as a ToxA interactor.

Despite the progress made in previous studies, the molecular basis underlying ToxA-induced necrosis and the ToxA-*Tsn1* interaction remain unknown. To better understand the molecular mechanism of ToxA during disease, we have re-analysed potential ToxA host interacting partners using a complementary series of protein-protein interaction approaches. We demonstrate that ToxA interacts in the apoplastic space with an integral membrane protein. In contrast, our reanalysis of previously identified ToxA-interacting proteins questions their validity as interacting partners and their role in ToxA-mediated disease. The identification of a valid interacting protein partner to the engimatic ToxA effector protein provides the basis to advance our understanding on how ToxA induces *Tsn1*-mediated cell death.

## Results

### ToxA binds to a wheat transmembrane protein TaNHL10

A yeast two-hybrid (Y2H) library screening approach was used to identify the host targets of ToxA. A prey library was constructed from *P. nodorum*-infected (at 2, 3, 4, and 5-dpi) and ToxA-infiltrated leaves (at 12, 24, 48, and 72-hpi) from the *Tsn1* wheat cultivar Grandin. The resulting Y2H library had 7.6 × 10^6^ independent clones and a cell density of 11.6 × 10^7^ cfu/mL against ToxA as bait. Initial screening of 5.9 × 10^6^ mated diploids resulted in over 300 colonies, which were further tested for production of yeast galactosidase (MEL1) resulting in 122 positive clones. These clones were subjected to colony PCR and restriction digest analysis to identify redundant inserts reducing the number of positive clones to 48, all of which were sequenced. Sequencing revealed 24 different cDNA clones from wheat including the clones of a partial cDNA of ToxABP1 (Manning *et al*., 2007), plastocyanin (Tai et al. 2007), and a pathogenesis-related protein 1 (PR-1) (Lu *et al*., 2014); all previously reported ToxA interactors. The plasmids of these clones were isolated and tested for prey auto-activity to eliminate false positives. Nineteen of these plasmids, including ToxABP1 and plastocyanin, showed auto-activity in the absence ToxA (Figure S1). No auto activity was detected for a pathogenesis related protein 1 (PR-1), an unidentified protein with FAF (fantastic four) domain or a predicted NDR1/HIN1-like (NHL) protein confirming these as putative ToxA-interacting proteins.

The NHL proteins belong to a large gene family in *Arabidopsis thaliana* (Zheng *et al*., 2004) and several are reported to be involved in plant immunity, including pathogen perception (Zipfel *et al*., 2004; Zheng *et al*., 2004; Boudsocq *et al*., 2010) and disease resistance (Aarts *et al*., 1998; Century *et al*., 1995). In addition, previous studies showed that NHL proteins are membrane associated and hypothesized as intercellular transducers of pathogen signals (Coppinger *et al*., 2004; Selote *et al*., 2014). Furthermore, previous studies have suggested that ToxA may interact with an extracellular receptor-like protein via its predicted RGD cell attachment motif (Meinhardt *et al*., 2002; Sarma *et al*., 2005; Manning *et al*., 2008), which is thought responsible for its internalization into host cell. Collectively, these previous studies have implicated a potential role for NHL proteins in ToxA-mediated disease and we therefore focused on the NHL protein (here after named TaNHL10) for the remainder of this study.

The identified TaNHL10 prey plasmid encodes a partial protein missing the first 93 amino acids (TaNHL10^94-210^). Neither the ToxA nor TaNHL10^94-210^ constructs showed auto-activity of the reporter genes when expressed with empty vector controls (Figure 1a), and protein expression was confirmed by immunoblotting (Figure S2). Subsequent analysis with full-length TaNHL10 revealed that it was unable to interact with ToxA. This was not unexpected as TaNHL10 has a predicted transmembrane domain (21-43 aa, by TMHMM) which could prevent its availability in the intracellular environment, where the interaction occurs in classical Y2H (Brückner *et al*., 2009).

**Figure 1.**
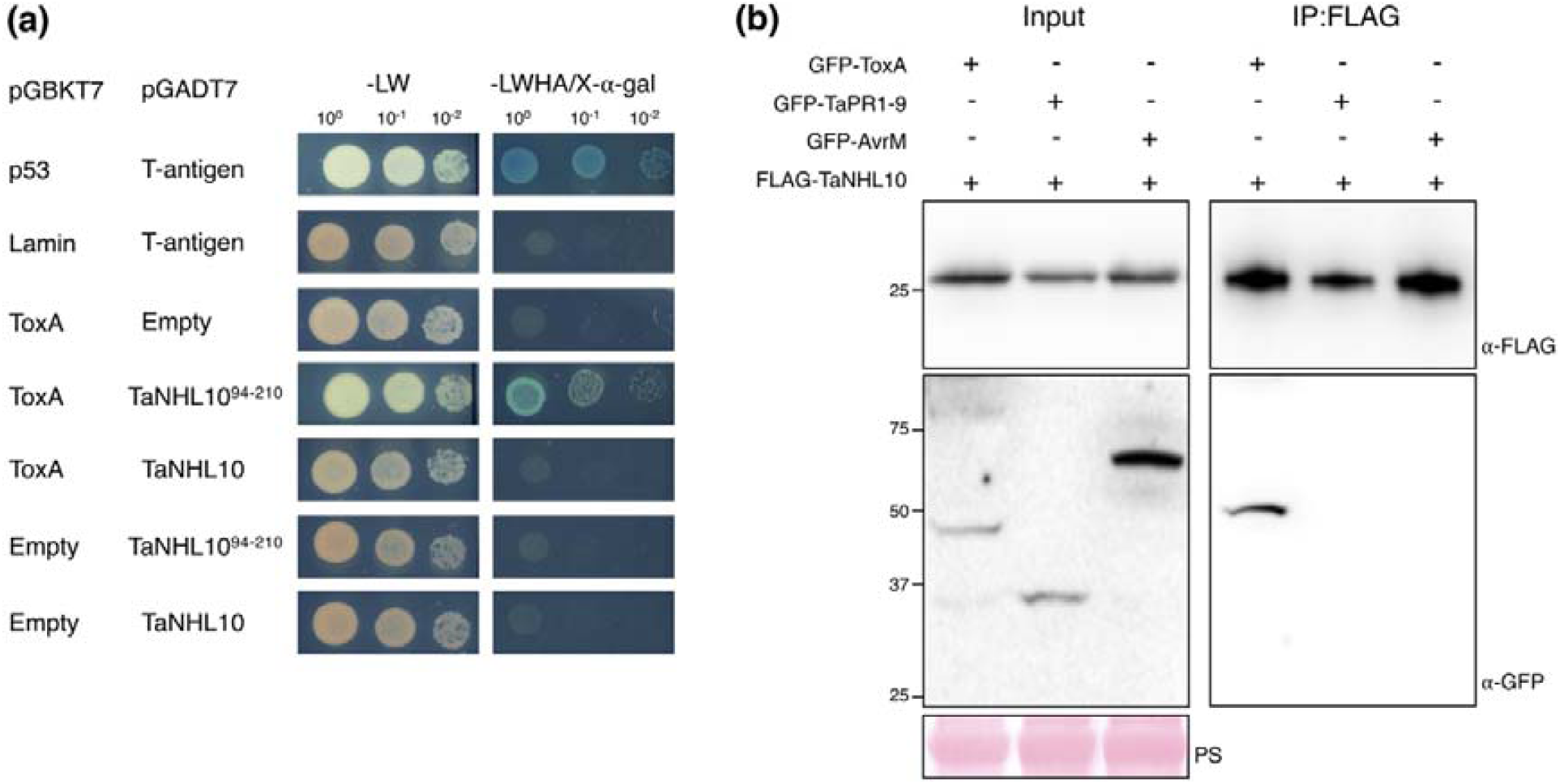
ToxA interacts with the wheat protein TaNHL10 in yeast-two-hybrid (Y2H) and *in planta* co-immunoprecipitation assays. (a) Y2H assay using ToxA (lacking signal peptide, 1-16 aa) and partial TaNHL10 (TaNHL10^94-210^) or full-length TaNHL10 proteins. Yeast co-expressing bait and prey plasmids were serially diluted from cell suspensions of a single yeast colony. Growth on -Leu,Trp (-LW) indicates the presence of both plasmids in the yeast colony. Growth and blue colouration on -Leu,Trp,His,Ade/X-⍰-Gal (-LWHA/X-⍰-Gal) is evidence of protein interaction. Serial dilutions reflect the strength of the interaction. p53/T-antigen and Lamin/T-antigen are positive and negative controls, respectively. ToxA was co-transformed with either empty vector, partial TaNHL10 (TaNHL10^94-210^), or full-length TaNHL10, in rows 3 to 5. TaNHL10^94-210^ and TaNHL10 were also co-transformed with an empty vector in the final two rows. **(b)** *in planta* co-immunoprecipitation between ToxA and TaNHL10. FLAG-TaNHL10 (25 kDa) was transiently co-expressed with either GFP-ToxA (45 kDa), GFP-TaPR1-9 (44 kDa), or GFP-AvrM (61 kDa) in *N. benthamiana*. Total protein extracts were incubated with Anti-FLAG beads. The captured proteins were immunoblotted. Ponceau S (PS) stain shows protein loading. Numbers on the left indicate molecular weight marker.

### Confirmation of the ToxA-TaNHL10 interaction *in planta*

To validate the ToxA-TaNHL10 interaction, we conducted *in planta* co-immunoprecipitation (Co-IP) experiments using the model plant system *Nicotiana benthamiana*. To do this, we fused GFP to the N-terminus of mature ToxA (GFP-ToxA) and a FLAG peptide to the N-terminus of full-length TaNHL10 (FLAG-TaNHL10), which were transiently co-expressed in leaves of *N. benthamiana* by agroinfiltration. Wheat pathogenesis-related protein 1-9 (GFP-TaPR1-9) and the flax-rust effector AvrM (GFP-AvrM) were used as controls. The proteins were extracted from the leaves two days after infiltration and immunoprecipitated using Anti-FLAG beads. GFP-ToxA was co-purified with the FLAG-TaNHL10 protein complexes; in contrast, there was no evidence of GFP-TaPR1-9 or GFP-AvrM co-purification (Figure 1b). Collectively, Y2H and *in planta* Co-IP experiments suggest that ToxA interacts with the host protein TaNHL10.

### ToxA specifically interacts with the LEA2 domain of TaNHL10

The TaNHL10 protein sequence includes a predicted transmembrane domain (21-43 aa) and a Late Embryogenesis Abundant 2 (LEA2) domain (85-174 aa) (Figure 2a). Our Y2H and *in planta* Co-IP data demonstrate that ToxA interacts with the partial sequence of TaNHL10 (TaNHL10^94-210^). To resolve where ToxA interacts in the C-terminal TaNHL10 region, we tested the interaction with the LEA2 domain (TaNHL10^94-174^) and the C-terminus of the protein (TaNHL10^175-210^). In addition, we also generated TaNHL10^1-20^ and TaNHL10^44-210^, where we excluded the predicted transmembrane domain (21-43 aa). There was no growth of the yeast strains harbouring TaNHL10^1-20^, TaNHL10^44-210^ and TaNHL10^175-210^ suggesting that these proteins did not interact with ToxA (Figure 2a), whereas the construct TaNHL10^94-174^ showed interaction with ToxA. The TaNHL10^94-174^ construct showed no auto-activity when co-transformed with empty vector control (Figure 2b), and protein expression was confirmed by immunoblotting (Figure S2).

**Figure 2.**
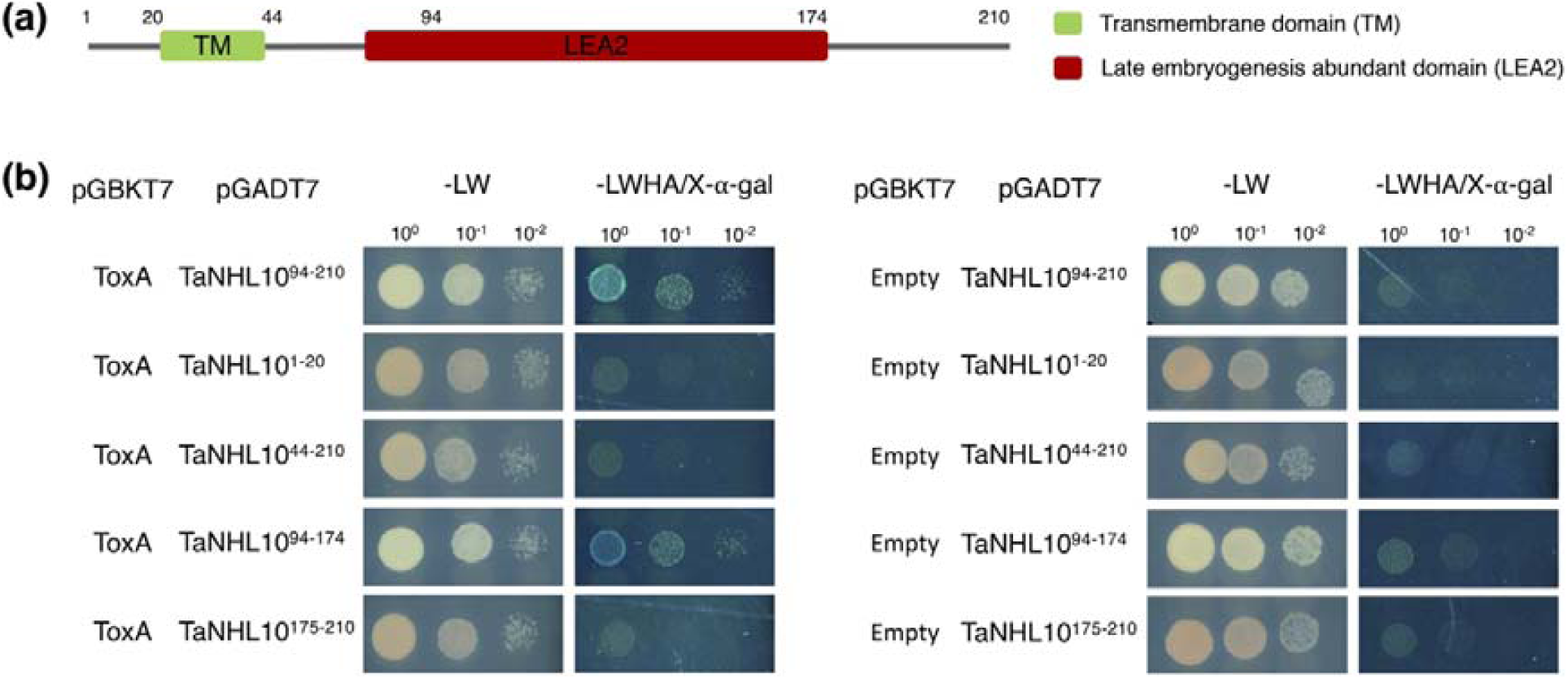
ToxA specifically interacts with the LEA2 domain of TaNHL10. **(a)** Schematic view of TaNHL10 domain organization. The transmembrane (TM) domain is shown in green and C-terminal Late embryogenesis abundant domain (LEA2) is shown in red. **(b)** Y2H confirmation assay. ToxA was co-transformed with either empty vector or one of the TaNHL10 truncate constructs (TaNHL10^1-20^, TaNHL10^44-210^, TaNHL10^94-174^ or TaNHL10^175- 210^). Yeast co-expressing bait and prey plasmids were serially diluted from cell suspensions of a single yeast colony. Growth on -LW indicates the presence of both plasmids in the yeast colony, and growth and blue coloration on -LWHA/X-⍰-Gal shows the interaction of the two proteins. Serial dilutions reflect the strength of the interaction. TaNHL10^94-210^ served as a positive control.

A bioinformatic search using the genome of wheat cultivar Chinese Spring revealed five homologous genes with high sequence similarities to TaNHL10. As we previously identified interaction between TaNHL10^94-174^ (the partial LEA2 domain of TaNHL10) and ToxA, we tested this region from the TaNHL10 homologs against ToxA. The results from our Y2H assay showed that this region from all five homologous proteins also interacted with ToxA (Figure S3), indicating that ToxA can interact with all identified TaNHL10 homologs in wheat.

### Membrane topology of TaNHL10

Next, we were interested in understanding the cellular location of the ToxA-TaNHL10 interaction. Based on the transmembrane area prediction software, TMHMM2.0 (Krogh *et al*., 2001; Sonnhammer *et al*., 1998), we speculated that the C-terminal LEA2 domain of TaNHL10 (Figure 2a) was extracellular. To experimentally validate this, we performed a subcellular co-localization analysis of N-terminally GFP fused TaNHL10 (GFP-TaNHL10) using the established transmembrane marker Remorin from *Solanum tuberosum* (StREM1.3) in *N. benthamiana* (Raffaele *et al*., 2009; Reymond *et al*., 1996). Confocal microscopy analysis demonstrated that the GFP-TaNHL10 accumulated mainly at the cell periphery, and that the GFP-TaNHL10 signal almost completely overlapped with the fluorescence signal from RFP-labelled StREM1.3 (RFP-REM1.3) (Figure 3b, Figure S4a). In contrast, analysis of the GFP control showed localization to the nucleus and cytoplasm (Figure 3a). The fluorescence signal of GFP-TaNHL10 remained associated with the plasma membrane after plasmolysis of the cells (Figure S5). Similar membrane co-localization was observed with TaNHL10 fused to RFP at the N terminus (RFP-TaNHL10) and GFP-labelled StREM1.3 (GFP-REM1.3) (Figure 3c). However, when we expressed C-terminally RFP-labelled TaNHL10 (TaNHL10-RFP), the RFP fluorescence signal did not co-localize with GFP-REM1.3 but was observed between GFP-REM1.3 membrane signals from neighbouring cells (Figure 3d), suggesting the RFP signal in the apoplast. Western blot analysis revealed that the RFP-TaNHL10 samples showed an expected full-length (51 kD) protein band (Figure S4b), while TaNHL10-RFP samples showed a band size of 25-27 kD (Figure S4b), consistent to the RFP alone size. We also observed a similar-sized band in samples expressing C-terminally RFP-labelled secreted-ToxA (NtPR1sp-ToxA-RFP) (Figure S4b), suggesting the protein tag (RFP; 27 kD) was cleaved in the apoplast as previously reported (Van Esse *et al*., 2006; Di *et al*., 2016; Mesarich *et al*., 2016; Doehlemann *et al*., 2009). Together, these data reveal that TaNHL10 is membrane localized, and has a short intracellular N-terminus (predicted as 1-20 aa) with the remainder of the protein localized outside the cell (predicted as 44-210 aa).

**Figure 3.**
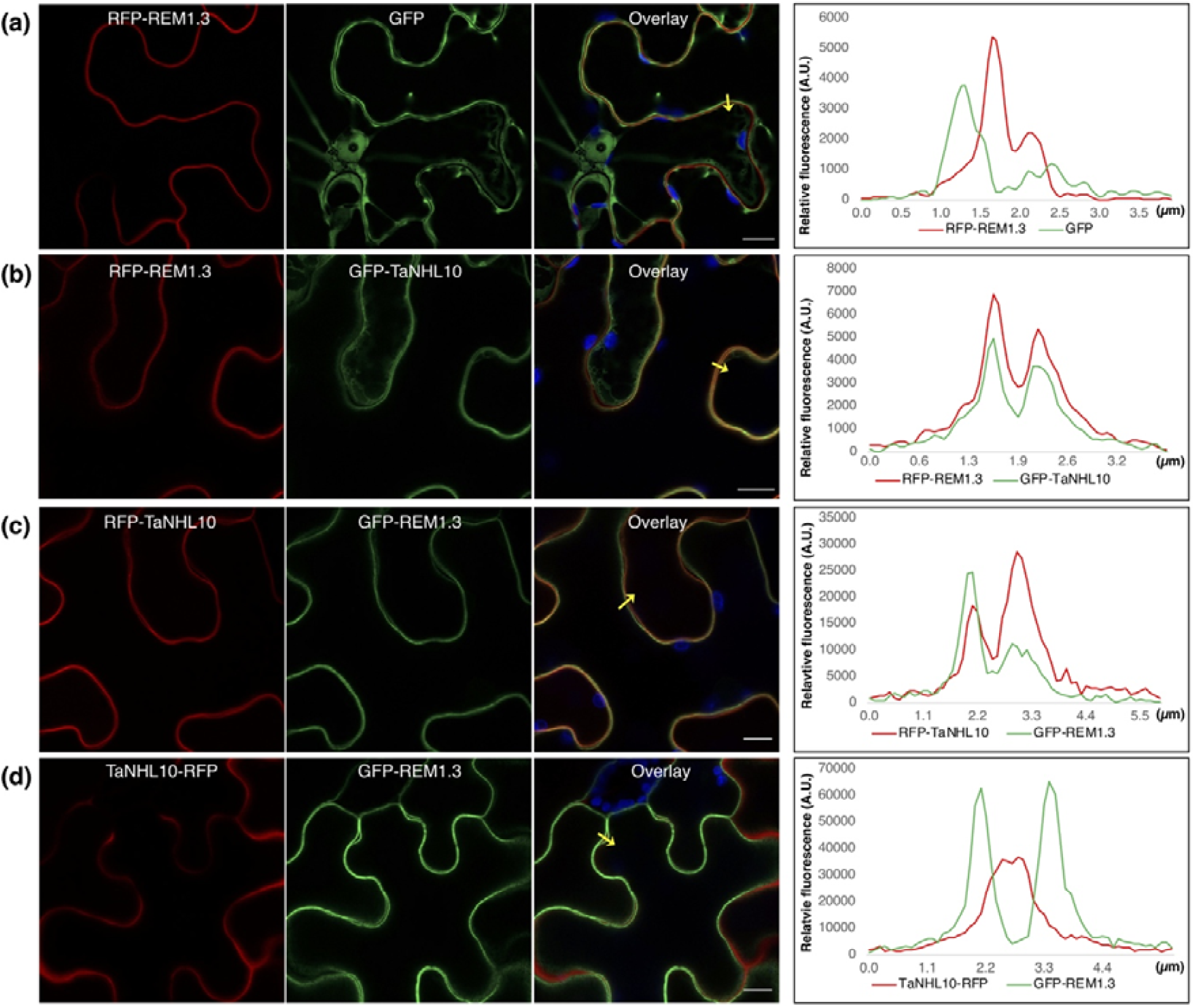
TaNHL10 localizes to the plasma membrane, and its C-terminus is extracellular *in planta*. *N. benthamiana* plants were transiently co-expressed by agroinfiltration using the following constructs: **(a)** RFP-REM1.3 (pK7WGR2/StREM1.3), as plasma membrane marker, and GFP (pK7WGF2-EV), as nucleo-cytoplasmic marker; **(b)** RFP-REM1.3 and GFP-NHL10 (pK7FWG2/TaNHL10); **(c)** RFP-NHL10 (pK7WGR2/TaNHL10) and GFP-REM1.3 (pB7WGF2/StREM1.3); and NHL10-RFP (pK7RWG2/TaNHL10) and GFP-REM1.3. Fluorescence intensities of RFP/GFP in membrane transects (yellow arrowheads) at 2-d post agroinfiltration. Chloroplast autofluorescence is in blue in overlay images. Scale bars represent 10 µm.

### Secreted ToxA co-localizes with TaNHL10

Complementary protein interaction approaches described above demonstrated that ToxA interacts with TaNHL10. However, due to the suspected cleavage of the linker between ToxA and RFP by apoplastic proteases (Figure S4b), we were unable to show the interaction using secreted ToxA by *in planta* Co-IPs. To mitigate this, we designed a new construct harbouring the *Nicotiana tobaccum* PR1 secretion signal (NtPR1sp), RFP with the reduced linker (-GSG-), and ToxA (NtPR1-RFP-ToxA) (Figure S6a), and transiently expressed the construct in *N. benthamiana*. Western blot analysis of protein expression in *N. benthamiana* showed the presence of the full-length recombinant protein and no evidence of the previously observed cleavage (Figure S4c and S6b). Consequently, we used this construct to address whether secreted ToxA associates with TaNHL10 *in planta* using confocal microscopy. Notably, in the presence of GFP-TaNHL10, the localization of secreted-RFP-ToxA shifted toward the cell membrane, resulting in a distinct overlap with fluorescence signals from the TaNHL10 (Figure 4a, Figure S7a). However, we observed no such shift in the signal in control leaf samples expressing GFP-REM1.3 (Figure 4b, Figure S7b) validating the ToxA-TaNHL10 interaction and indicating that ToxA associates with TaNHL10 at the plant cell surface.

**Figure 4.**
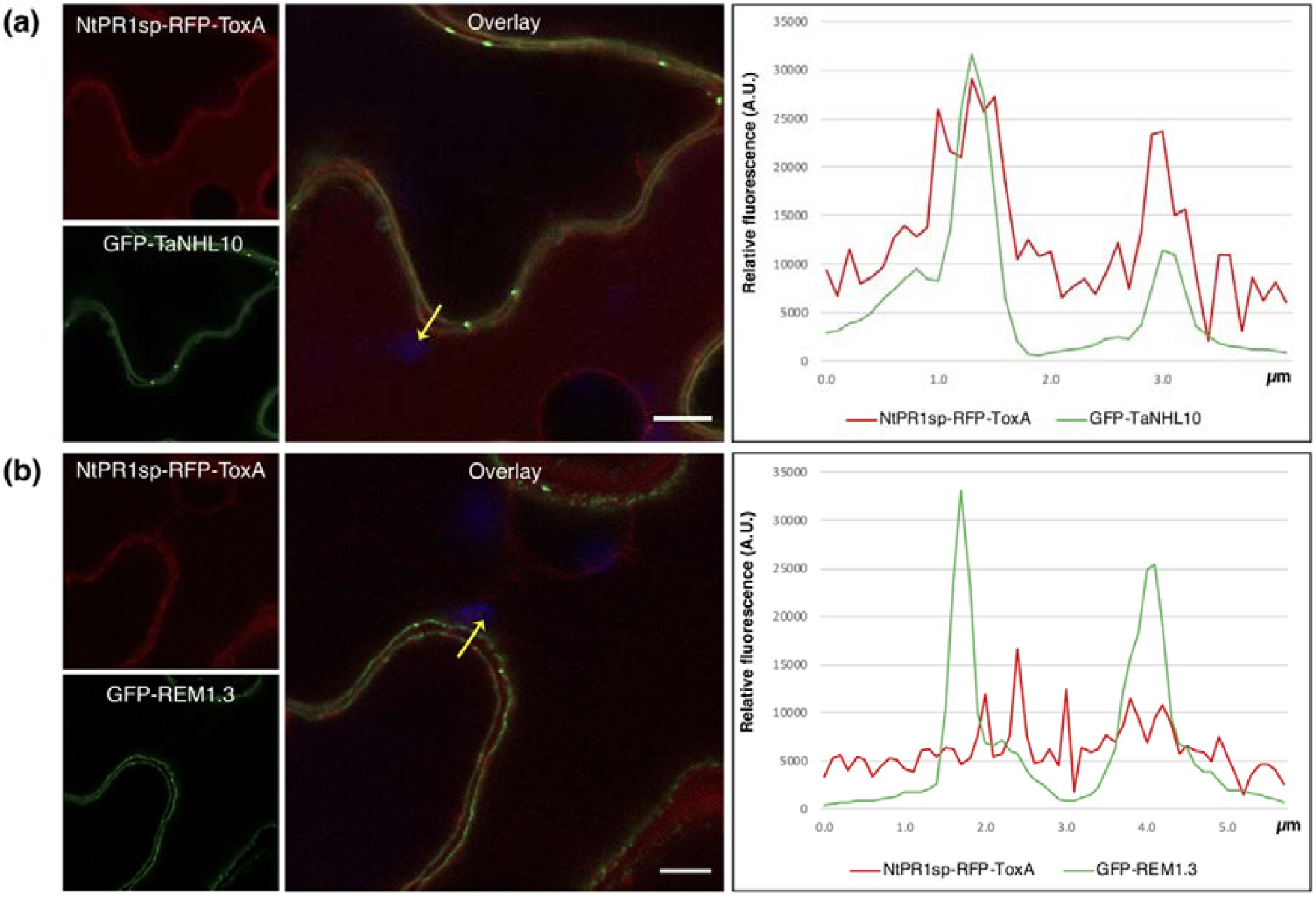
Secreted-ToxA co-localizes with TaNHL10 *in planta*. *N. benthamiana* plants were transiently co-expressed by agroinfiltration using the following constructs: **(a)** GFP-NHL10 and NtPR1sp-RFP-ToxA (pK7FWR2/NtPR1sp-mRFP-ToxA); and **(b)** GFP-REM1.3 and NtPR1sp-RFP-ToxA. The plasmolysis of the leaf cells was induced before imaging. Fluorescence intensities of RFP/GFP in membrane transects (yellow arrowheads) at 2-dpi. Chloroplast autofluorescence is in blue in overlay images. Scale bars represent 5 µm.

### *Tsn1*-mediated cell death is not active in the model plant systems

Our interaction studies have unequivocally demonstrated that ToxA and TaNHL10 interact. However, the functional characterization of the ToxA and TaNHL10 interaction in wheat would be slow and cumbersome due its lack of genetic amenability. As such, we attempted to recapitulate *Tsn1*-mediated cell death in model plant systems, such as *N. benthamiana* and *N. tobaccum*, which allow rapid functional assays. We co-expressed ToxA (with or without signal peptide), TaNHL10 and Tsn1 without any tags as well as various different recombinant versions of these three proteins in *N. benthamiana* and *Nicotiana tobaccum*. However, these attempts failed to produce cell-death in both model systems (data not shown). Furthermore, we expressed a predicted auto-active Tsn1 (Tsn1^D728V)^) (Bendahmane *et al*., 2002; Howles *et al*., 2005; Van Ooijen *et al*., 2008; Adachi *et al*., 2019) in both plant systems, which also did not result in visible cell death (Figure S8a). We were able to detect protein expression of full-length Tsn1 and its auto-active mutant in *N. benthamiana* (Figure S8b). Altogether, this indicates that these model plants are likely lacking the downstream component(s) required for *Tsn1*-mediated cell death.

### The ToxA-TaNHL10 interaction is required for ToxA-mediated necrosis

To determine the biological relevance of ToxA and TaNHL10 interaction, we sought to identify mutations in ToxA that abolished interaction with TaNHL10 and assess the ability of these non-interacting mutants to trigger necrosis in *Tsn1* wheat cultivars. A random PCR mutagenesis approach was used to generate a library of mutant ToxA genes that was screened for loss of interaction with TaNHL10 using a Y2H approach. Amino acid mutations at position 109 (ToxA^N109D^) and 84 (ToxA^I84T^) (Figure 5a) abolished binding of ToxA with TaNHL10. Subsequent immunoblotting analysis showed that ToxA wild-type, ToxA^N109D^, and ToxA^I84T^ proteins were expressed at comparable levels in yeast (Figure S9). Using the known structure of ToxA (PDB ID: 1ZLD) (Sarma *et al*., 2005), we verified that N109 is surface-exposed (Figure 5b), whilst I84 was buried in the structure (Figure 5b).

**Figure 5.**
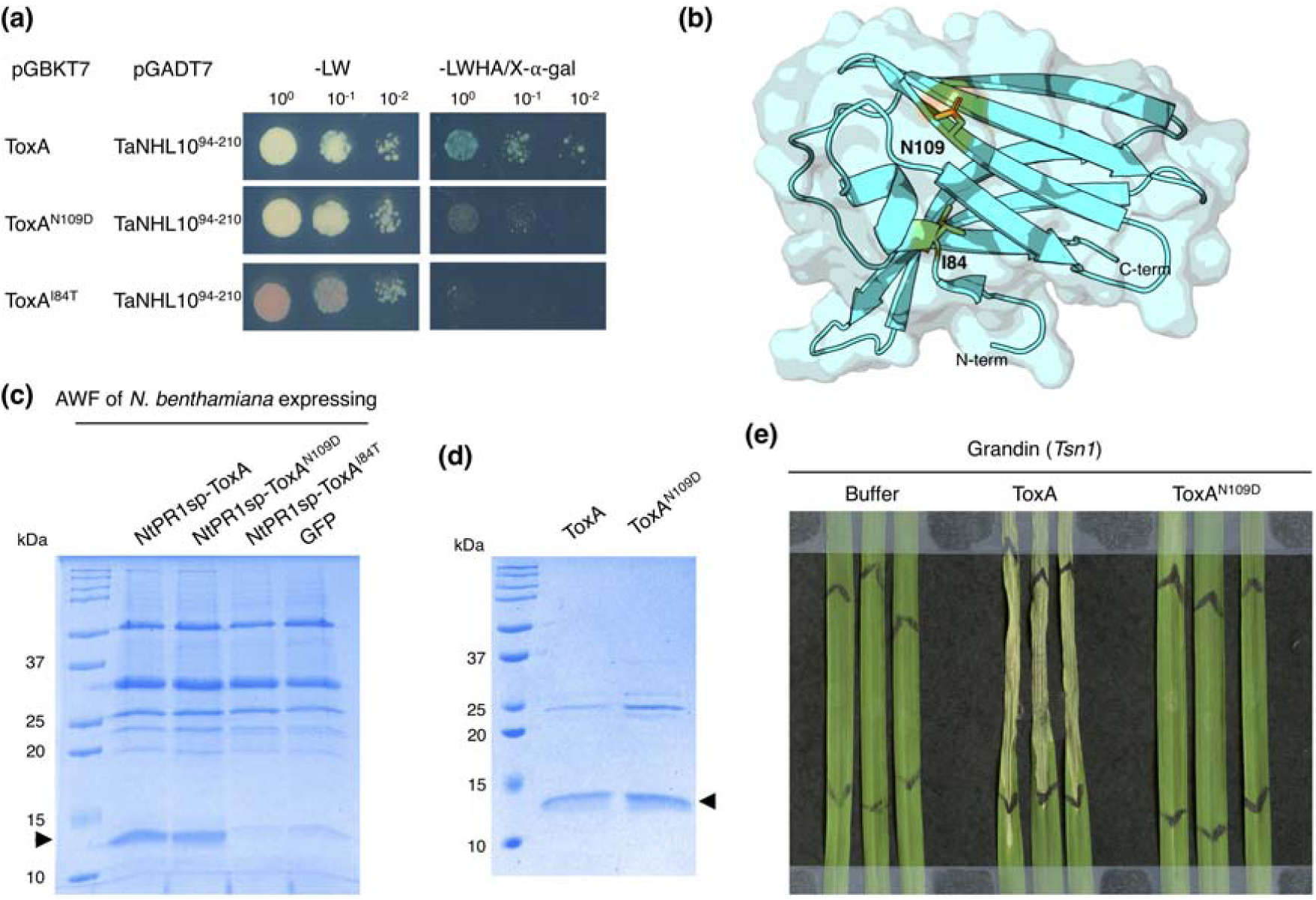
A single point mutation in ToxA abolishes interaction with TaNHL10 as well as recognition by *Tsn1*. **(a)** Yeast-two-hybrid assays using ToxA, ToxA^N109D^ and ToxA^I84T^ as baits, and TaNHL10 as prey. Yeast co-expressing bait (pGBKT7) and prey (pGADT7) plasmids were serially diluted from cell suspensions of a single yeast colony. Growth on -Leu,Trp (-LW) indicates the presence of both plasmids in the yeast colony. Growth and blue coloration on -Leu,Trp,His,Ade/X-⍰-Gal (-LWHA/X-⍰-Gal) show the interaction of two proteins. Serial dilutions reflected the strength of the interaction. **(b)** Cartoon and surface representation of ToxA (PBD ID: 1ZLD). Residues I84 and N109 are shown in stick representation and coloured. **(c)** ToxA and its variants’ expression in *Nicotiana benthamiana*. Coomassie Brilliant blue stained sodium dodecylsulfate-polyacrylamide gel electrophoresis (SDS-PAGE) analysis of apoplast washing fluid (AWF) isolated from *N. benthamiana* expressing various constructs. The black arrow indicates the expected size of the matured ToxA and its variants (13.8 kDa). AWF from *N. benthamiana* expressing GFP was used for background control. **(d)** Coomassie-stained SDS-PAGE showing amount comparison of ToxA and ToxA^N109D^ after size-exclusion chromatography of AWFs from *N. benthamiana*. The black arrow indicates the expected size of the mature ToxA and ToxA^N109D^ proteins (13.8 kDa). **(e)** Syringe-infiltration of buffer control (10mM HEPES pH 7.5, 150 mM NaCl), ToxA, and ToxA^N109D^ into the second leaf of 2-week-old Grandin (*Tsn1*). The black markings on the leaves indicate the infiltration zone. Leaves were harvested 3-day post infiltration and imaged.

To assess if the mutants were able to induce *Tsn1*-mediated cell death, ToxA, ToxA^N109D^, and ToxA^I84T^ were produced as secreted forms in *N. benthamiana* and isolated from the apoplast washing fluid (AWF) (Dagvadorj and Solomon, 2021). The AWF with secreted ToxA and ToxA^N109D^ showed bands representing ToxA (between 10-15 kDa) of similar size and intensity on SDS-PAGE analysis (Figure 5c), whereas this band was absent in ToxA^I84T^ and GFP-expressing *N. benthamiana* AWFs (Figure 5c). Collectively, these data suggest that the internal structural changes in the ToxA^I84T^ mutant had an adverse impact on folding and/or protein stability preventing accumulation in apoplast. We subsequently removed ToxA^I84T^ from further analysis. The ToxA and ToxA^N109D^ proteins within the AWF were further purified from the apoplast by size-exclusion chromatography (SEC) (Figure 5d). The SEC chromatograms (Figure S10a) and SDS-PAGE analysis of the SEC fractions (Figure S10b) showed that the ToxA^N109D^ mutant eluted at the same time as the wild-type ToxA. We confirmed the integrity of both proteins using intact protein mass spectrometry under non-reducing conditions (Table S2 and Figure S11). Based on the mass obtained from these data, we confirmed that ToxA and ToxA^N109D^ were purified from the apoplast with the single disulfide bond intact, further confirming the integrity of the protein. Semi-purified preparations of both ToxA and Tox^N109D^ (Figure 5c) were infiltrated into the *Tsn1* wheat line Grandin to assess whether ToxA^N109D^ could induce necrosis at a comparable level to ToxA. As expected, strong necrosis was evident in the *Tsn1* wheat cultivar subsequent to ToxA infiltration (Figure 5e, Figure S12). However, necrosis was not observed in ToxA^N109D^ - infiltrated Grandin (Figure 5e, Figure S12), indicating that the interaction of ToxA with TaNHL10 is required for *Tsn1*-mediated necrosis.

## Discussion

Here, we sought to identify and characterise wheat proteins that interact with ToxA and facilitate *Tsn1*-mediated cell death. We identified that ToxA targets the transmembrane TaNHL10 protein using a yeast-two-hybrid (Y2H) screening approach. Consistent with previous reports, ToxABP1 (Manning *et al*., 2007), plastocyanin (Tai et al. 2007), and a pathogenesis-related protein 1 (PR-1) (Lu *et al*., 2014) were all identified in our initial Y2H screening. Surprisingly, the subsequent bait dependency test revealed that ToxABP1 and plastocyanin clones are auto-active in the absence of ToxA (Figure S1). Whilst we did not assay full-length ToxABP1 or plastocyanin for auto-activity, the auto-active phenotypes observed for the partial clones raise some concerns regarding the previous published data and question the validity of these interactions. It should also be noted that, in these previous studies, the protein-protein interaction experiments were restricted to Y2H only (Manning *et al*., 2007; Tai *et al*., 2007). In this study, we have exploited complementary approaches to identify and confirm the direct interaction of ToxA with TaNHL10.

The *NHL* genes belong to a large family in *A. thaliana*. Within the NHL proteins, AtNDR1 from *A. thaliana* is arguably one of the most recognized and intensively studied. Several studies reported that NDR1 proteins from different species are localized in the plasma membrane (Coppinger *et al*., 2004; Selote *et al*., 2014; Yang *et al*., 2021; Liu *et al*., 2020). Because of the predicted topology of NDR1 in the plasma membrane, it has been thought that it is involved in extracellular activity as a signalling protein between the apoplast and plant cells (Coppinger *et al*., 2004). Further studies have supported this idea by showing that AtNDR1 is involved in plant cell wall adhesion (Knepper *et al*., 2011a). Hence, it has been hypothesized that the NDR1 proteins have a transmembrane domain, with the C-terminus exposed to the apoplast (Coppinger *et al*., 2004; Selote *et al*., 2014); however, this has not been confirmed experimentally. Our study is the first that provides experimental evidence of the membrane topology of an NHL protein. Our result supports the previous hypothesis on NDR1 proteins by showing that TaNHL10 contains a transmembrane domain at its N-terminus and an extracellular C-terminus.

What might the role be of surface exposed NHL proteins in plant immunity? Our data shows that the ToxA effector specifically interacts with the extracellular part of TaNHL10, revealing that the interaction is apoplastic and occurs at the plant cell surface. Although this is the first report of an effector (a pathogen-derived molecule) targeting the extracellular domain of an NHL protein, it is likely that many NHL proteins sense different pathogen molecules like effectors. There have also been overwhelmingly consistent data from previous studies that *NHL* genes play roles in enhancing plant disease resistance (Gopalan *et al*., 1996; Selote *et al*., 2014; Century *et al*., 1997; Liu *et al*., 2020). Thus, we can speculate that NHL proteins localized at the cell surface, close to the infection site, facilitate instant sensing of pathogens or pathogen derived molecules, thereby activating quick defence responses (Knepper *et al*., 2011b; Varet *et al*., 2003; Liu *et al*., 2020).

We demonstrated that the interaction between ToxA and TaNHL10 was required for *Tsn1*-mediated cell death via mutagenesis studies of the effector. While we were interested to perform analogous experiments with TaNHL10, these remain to be time consuming and challenging experiments in the wheat due to its complex and genetically intractable nature. Alternatively, we tried to observe *Tsn1*-mediated cell death in the model plant systems such as *N. benthamiana* and *N. tobaccum*, which would allow the analysis of the interactions between effector (ToxA) and its cognate receptor (Tsn1), as has been previously described for similar systems (Vleeshouwers *et al*., 2008; Van Der Hoorn *et al*., 2000; Ma *et al*., 2012; Chen *et al*., 2017). However, these possibilities were restricted as neither ToxA, TaNHL10 and Tsn1 co-expression or autoactive Tsn1 mutant expression produce visible cell-death in *N. benthamiana* or *N. tobaccum*. These outcomes prevented us from further characterising ToxA recognition by *Tsn1* in model plant systems.

So, how might TaNHL10 be involved in ToxA recognition by *Tsn1*? Manning et al. (2008) showed that the C-terminal RGD motif of ToxA is necessary for internalization of the effector into host plant cells, which is possibly regulated by the interaction between ToxA and an extracellular receptor via the RGD motif. We have demonstrated that TaNHL10 is an extracellular surface protein that associates with ToxA. However, mutation of the RGD motif in ToxA (ToxA^RGD^ to ToxA^AAA^) resulted in a comparable level of interaction with TaNHL10 to that of wild-type ToxA (Figure S13). Therefore, the RGD motif in ToxA is not required for interaction with TaNHL10. We cannot though exclude the possibility that TaNHL10 may be a surface receptor that mediates ToxA trafficking into plant cells using endocytosis. Future work is needed to investigate whether the delivery of ToxA into the plant cell is coupled with TaNHL10.

Previous studies have suggested that the NHL proteins could have several biological functions. For example, *Atndr* mutants fail to show resistance mediated by the CC-NLRs RPM1 and RPS2 to avirulent *Pseudomonas syringae* producing AvrRpm1, AvrB, and AvrPrt2 effectors (Century *et al*., 1995; Aarts *et al*., 1998), suggesting that AtNDR1 is a positive regulator involved in effector-triggered immunity (ETI). It has also been demonstrated that *Atndr* mutants show increased susceptibility to virulent *Pseudomonas syringae*, whereas *AtNDR1* overexpression promotes resistance against virulent *P. syringae* (Coppinger *et al*., 2004), indicating that in addition to its role in ETI, AtNDR1 functions in PAMP-triggered immunity (PTI). Furthermore, a more recent study revealed that *Atndr1* mutant plants displayed electrolyte leakage and altered plasma membrane and cell wall adhesion (Knepper *et al*., 2011a), thereby indicating that AtNDR1 also has roles plant homeostasis. Our study has demonstrated that ToxA requires interaction with TaNHL10 to induce *Tsn1*-mediated necrosis. It is relevant to note that the Tsn1 bears a striking resemblance to typical NLR resistance proteins (Faris *et al*., 2010), and it has been previously proposed that necrotrophic pathogens such as *P. nodorum* may hijack host resistance proteins for their own benefit (Faris *et al*., 2010; Shi *et al*., 2016; Faris and Friesen, 2020). Therefore, as AtNDR1 is involved in ETI, we speculate that TaNHL10 is an important component in *Tsn1*-mediated cell death pathways, and is manipulated by ToxA, ultimately promoting the lifecycle of the necrotrophic pathogen. The next steps in this research are to determine the molecular mechanism of TaNHL10 and understand its interplay between ToxA and Tsn1. These data advance our current understanding on how these pathogens manipulate host plants to promote disease and provide a platform for future research focussed on understanding the interplay between ToxA, Tsn1 and TaNHL10.

### Experimental procedures

#### Plants and fungal growth conditions

Wheat (*Triticum aestivum*) lines Grandin (*Tsn1*) and Corack (*tsn1*) were cultivated in a plant growth chamber with 250 μE light intensity, 85% relative humidity, and photoperiod of 16-h light at 20°C/8-h dark at 12°C. *Nicotiana benthamiana and Nicotiana tobaccum* plants were grown at 22°C with 16-h light/8-h dark cycle in a plant growth room. *Parastagonospora nodorum* SN15 strain was cultured on V8-PDA medium at 22°C with 12-h light/12-h night cycle.

#### Cloning

For Y2H assays, a gBlock of *P. nodorum ToxA* lacking the signal peptide (17-178 aa, designated as *ToxA*) (Friesen *et al*., 2006) and full-length *TaNHL10* (amplified including its stop codon from Grandin genomic DNA) were cloned into pENTR/D-TOPO (pENTR-TaNHL10), then recombined into gateway compatible pGADT7-GW (pGADT7-TaNHL10) (Lu *et al*., 2010). Truncated DNA sequences of TaNHL10 (TaNHL10^1-20^, TaNHL10^44-210^, TaNHL10^94-174,^ and TaNHL10^175-210^) were amplified from pGADT7-TaNHL10, and digested with EcoRI and BamHI and cloned into pGADT7. For *in planta* co-immunoprecipitation experiments, pENTR-TaNHL10 was recombined into pEarlyGate202 for FLAG-TaNHL10 expression (Earley *et al*., 2006). To generate GFP-ToxA construct, ToxA was amplified and cloned into the pENTR/ D-TOPO vector, subsequently recombined into pB7WGF2.0 (Karimi *et al*., 2002) by gateway LR reaction. Likewise, GFP-TaPR1-9 construct was prepared using the same approach. For GFP-AvrM, the pENTR-AvrM-A vector from the previous study (Catanzariti *et al*., 2010) was recombined into pB7WGF2.0 vector. For confocal co-localization experiments, an RFP-TaNHL10 construct was generated by subcloning pENTR-TaNHL10 plasmids into pK7WGR2 (Karimi *et al*., 2002). To generate TaNHL10-RFP construct, the full-length TaNHL10 gene was amplified without stop codon (TaNHL10-ST) from Grandin wheat genomic DNA and cloned in pENTR/D-TOPO (pENTR-TaNHL10-ST) and was recombined into pK7RWG2 (Karimi *et al*., 2002). For GFP-REM1.3 and RFP-REM1.3 constructs, gBlock sequence of StREM1.3 (Genbank ID: U7289.1) (Reymond *et al*., 1996) was cloned into pENTR-D/TOPO and then sub-cloned into pB7WGF2 and pK7WGR2, respectively. pB7WGF2_empty vector was used for GFP expression. For secreted protein expression in *N. benthamiana*, the vector pGWB411-NtPR1sp-ToxA expressing secreted ToxA from Dagvadorj and Solomon (2021) was used. The secreted ToxA^N109D^ mutant was generated using the QuikChange Lightning Site-Directed Mutagenesis Kit (Agilent Technologies) following the manufacturer’s protocol. Single nucleotide mutation of ToxA (ToxA^N109D^) was generated by amplifying pGWB411-NtPR1sp-ToxA with primer pairs carrying the desired mutation. All the plasmids used in this research were transformed into *Escherichia coli* (NEB^®^ 5-alpha), and the sequences were confirmed by Sanger sequencing before transformation into the *Saccharomyces cerevisiae* strain Y2HGold (Clontech) for Y2H assays or *Agrobacterium tumefaciens* (GV3101, pMP90) for agroinfiltration assays.

#### Yeast-two-hybrid library construction and screen

A yeast-two-hybrid library specific for ToxA screening was constructed using the Make Your Own ‘Mate & Plate’ Library System (Clontech) following the manufacturer’s instructions. ToxA lacking signal peptide was heterologously expressed in *E. coli* and purified as previously described (Outram *et al*., 2021).

*T. aestivum* cultivar Grandin was either infiltrated with the purified ToxA or infected with *P. nodorum* strain SN15 as described previously (Tan *et al*., 2008). Leaf samples were collected at 12, 24, 48, and 72 hours post-infiltration with purified ToxA, or 2, 3, 4, and 5 days post-infection. Total RNA was extracted from the samples, and cDNA synthesis (Make Your Own ‘Mate & Plate’ Library System User Manual; Clontech) was carried out as described in the manufacturer’s instruction. Combined double-stranded cDNA samples (each containing ≈ 300 ng DNA) were used to generate the prey library according to the manufacturer’s recommendations (Make Your Own ‘Mate & Plate’ Library System User Manual; Clontech). For yeast mating, the bait construct pGBKT7-ToxA was transformed into Y2HGold (Clontech) and the prey library constructs in pGADT7-Rec (Clontech) into yeast strain Y187 (Clontech). Mated diploid colonies were screened for protein-protein interactions on medium without -L, -W, -H, and -A amino acids for growth and with X-α-gal for blue coloration. To identify mutations in ToxA that inhibited the binding with TaNHL10, PCR-based random mutagenesis of the ToxA gene (without the signal peptide) was performed using GeneMorph II Random Mutagenesis Kit (Agilent) following the manufacturer’s protocol. The PCR product, containing random mutations of ToxA, was co-transformed with the bait vector pGBKT7 (linearized with EcoRI and BamHI restriction enzymes) and pGADT7-TaNHL10^94- 210^ into Y2HGold competent cells and screened for a loss of interaction with TaNHL10^94-210^. The mutant ToxA^N109D^ was identified, and an additional Y2H confirmation test was performed to determine the interaction between ToxA or ToxA^N109D^ through SD/-L/-W/-H/-A/X-α-gal selection medium.

#### Co-immunoprecipitation experiments and immunoblot analysis

Proteins were transiently expressed by agroinfiltration in *N. benthamiana* as previously described (Dagvadorj *et al*., 2017), and the leaf samples were harvested at 2-3 day-post-agroinfiltration. Protein extraction, purification and western blot analysis were performed with minor modifications as described in Breen *et al*. (2016). Briefly, leaf samples were ground in liquid nitrogen and dissolved in extraction buffer GTEN containing 1 % (w/v) polyvinylpolypyrrolidone, 0.1 % Nonidet P-40 substitute (Amresco), 1 mM PMSF (Amresco), and 1 x protease inhibitor cocktail tablet EDTA-free (Roche). The cell debris was pelleted by centrifugation and the supernatant was used for co-immunoprecipitation (Co-IP). Anti-FLAG magnetic beads (Sigma) were used to immunoprecipitate FLAG fusion proteins.

Yeast total protein was extracted as described previously (Kushnirov, 2000). For western blot analysis, anti-myc (Santa Cruz Biotech), anti-FLAG (Thermo) and anti-GFP (Santa Cruz Biotech) from rabbit were used as the primary antibodies; and anti-rabbit HRP conjugated antibody (Sigma) was used as the secondary antibody. The anti-HA-Peroxidase from rat (Roche), and HRP conjugated anti-RFP from rabbit (abcam) were used to detect HA, and RFP-fusion proteins, respectively.

#### Confocal microscopy

Leaf discs were collected from *N. benthamiana* leaves 2-3 day-post-agroinfiltration, immersed in distilled water and imaged on Zeiss LSM 800 Airyscan confocal microscope. The excitation of GFP and RFP probes were performed by 488 and 561 nm laser diodes, respectively, and fluorescent emissions were detected at 495-550 nm (GFP) and 570-620 nm (RFP). Chloroplast autofluorescence was detected at 680-730nm. The GFP and RFP spectra were checked on experimental samples (GFP-NHL10 and RFP-REM1.3) on Zeiss 780 confocal microscope by Spectral Unmixing of Lambda mode data, confirming the residual channels, representing fluorescence emission from uncharacterized sources, was of low intensity. To induce plasmolysis, leaf samples were incubated in 0.45 M mannitol for 30 min before imaging. The images were processed by using ImageJ (Fiji) Software.

#### Apoplastic washing fluid isolation and size-exclusion chromatography

Apoplast washing fluid (AWF) was isolated from the *N. benthamiana* leaves expressing secreted ToxA and ToxA^N109D^ as previously described (Dagvadorj and Solomon, 2021). The AWFs with ToxA and ToxA^N109D^ proteins were further purified using size exclusion chromatography (SEC). For SEC, a S75 increase (10/300) was equilibrated with 10 mM HEPES pH 7.5 and150 mM NaCl, and 1 mL fractions were collected. Fractions were visualized by Coomassie-stained SDS-PAGE and those containing the protein of interest were collected for further analysis.

#### Intact mass spectometry (MS)

For intact protein MS analysis, 35 µL of collected fractions from SEC containing ToxA or ToxA^N109D^ proteins were adjusted to 5 μM in 0.1% formic acid (FA) for HPLC-MS analysis. The samples were then injected onto an Agilent UHPLC system. Each sample was first desalted for 2 min on an Agilent (Santa Clara, CA, USA) C3 trap column (ZORBAX StableBond C3) at a flow rate of 500 µl/min using a buffer containing 0.1% FA, followed by separation over 8 min using a gradient of 5–80% in a buffer containing 80% ACN/0.1% FA (ACN: acetonitrile), at a flow rate of 500 µl/min. The eluted material was directly analysed using an Orbitrap Fusion™ Tribrid™ mass spectrometer. MS acquisition was performed using the Intact Protein Mode script. The acquisition was performed across m/z 200-4000 with an accumulation time of 1 second. Data were analysed using the Free Style (v.1.4, ThermoFisher) protein reconstruct tool across a mass range of *m/z* 500– 2000, and protein masses between 10 and 20 kDa were searched.

#### AWF and SEC fraction infiltration into wheat leaves

AWF samples and SEC fractions containing ToxA and ToxA^N109D^ were infiltrated using a needleless syringe into the second leaf of 2-week old wheat cultivars until the infiltration zone reached 3-5 cm. The wheat leaves were checked for the induction of necrosis from 1 day post-infiltration (dpi) and recorded at 3 dpi.

### Accession numbers

*ToxA* (Genbank: DQ423483), *TaNHL10* (EnsemblPlants: TraesCS2B02G603100), *TaPR1-9* (EnsemblPlants: TraesCS5B02G442600), *StREM1*.*3* (GenBank: NM_001288060), *Tsn1* (Genbank: GU259634). All other relevant information can be found within the manuscript and its supporting materials.

## Acknowledgements

This research was supported by the Australian Research Council (ARC) Discovery Project DP180102355. The authors would like to acknowledge the facilities, and the scientific and technical assistance of Microscopy Australia at the Advanced Imaging Precinct, Australian National University. The authors thank Erin Hill for producing and purifying the ToxA effector that was used for Y2H library sample preparation. The authors also thank Daniel Yu for his technical assistance with the MS experiment.

## Short legends for supporting information

**Figure S1**. The ToxABP1 and plastocyanin clones showed autoactivity in the absence of ToxA.

**Figure S2**. Confirmation of protein expression in yeast.

**Figure S3**. Y2H assay screening ToxA interaction with five TaNHL10 homologs found in wheat genome.

**Figure S4**. Confirmation of protein expression in *N. benthamiana*.

**Figure S5**. TaHL10 localizes to the plasma membrane.

**Figure S6**. Short linker prevents the cleavage of RFP tag from RFP-ToxA in apoplast.

**Figure S7**. Secreted-ToxA co-localizes with TaNHL10 *in planta*.

**Figure S8**. The autoactive mutant of Tsn1 protein is not active in model plant systems.

**Figure S9**. A single point mutation in ToxA abolishes interaction with TaNHL10 as well as recognition by *Tsn1*.

**Figure S10**. Size exclusion chromatography of AWFs from *N. benthamiana* producing ToxA and ToxA^N109D^ proteins.

**Figure S11**. Intact ToxA and ToxA^N109D^ mass spectrometry analysis.

**Figure S12**. Genotype specificity of ToxA and ToxA^N109D^ samples.

**Figure S13**. The ToxA RGD motif is not required for ToxA-TaNHL10 interaction.

