## Supplementary data for "The necrotrophic effector ToxA from *Parastagonospora nodorum* interacts with wheat NHL proteins to facilitate *Tsn1*-mediated necrosis"

**Supplementary figures**

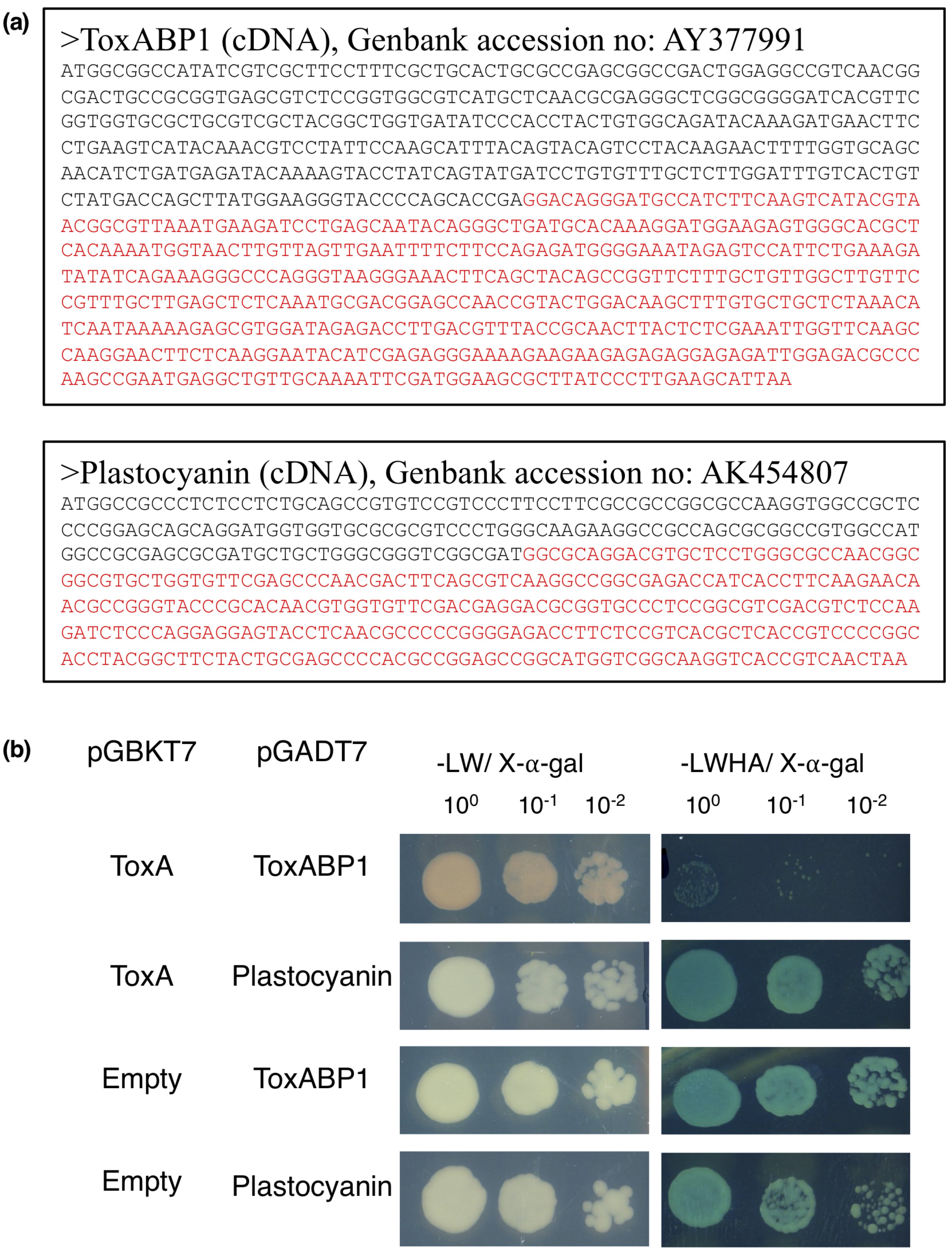

**Figure S1. The ToxABP1 and plastocyanin clones showed autoactivity in the absence of ToxA.** **(a)** The full-length cDNA sequences of ToxABP1 and plastocyanin and the sequence coverage of ToxABP1 and plastocyanin clones (in red) identified by Y2H screening against ToxA bait. **(b)** Yeast two-hybrid assay for testing auto-activity for ToxABP1 and plastocyanin clones identified Y2H screening against ToxA. Yeast co-expressing bait (pGBKT7) and prey (pGADT7) plasmids were serially diluted from cell suspensions of a single yeast colony. Growth on -Leu,Trp (-LW) indicates the presence of both plasmids in the yeast colony. Growth and blue coloration on -Leu,Trp,His,Ade/X-⍺-Gal (-LWHA/X-⍺-Gal) show the interaction of two proteins. Serial dilutions reflected the strength of the interaction. ToxABP1 and plastocyanin clones co-transfomed with empty bait vector showed auto-activity.

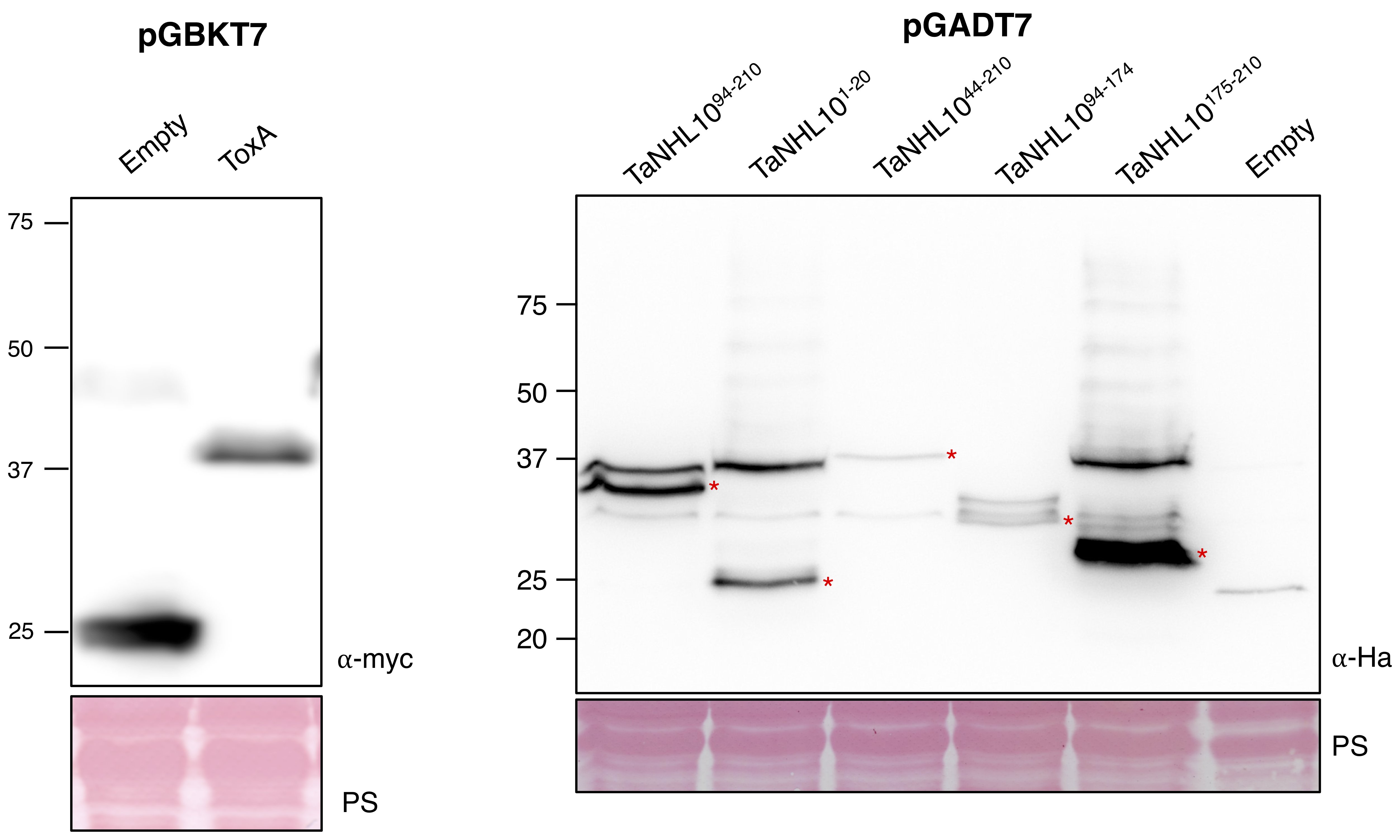

**Figure S2. Confirmation of protein expression in yeast.** ToxA (40.3 kDa) and TaNHL10 truncates, including TaNHL10^94-210^ (31.1 kDa), TaNHL10^1-20^ (20.6 kDa), TaNHL10^44-210^ (36.6 kDa), TaNHL10^94-174^ (27.4 kDa) and TaNHL10^175-210^ (22.3 kDa), protein expression in yeast. ToxA was detected with ⍺-myc and TaNHL10 truncate proteins were detected with ⍺-ha. Asterisks indicate the expected size of the recombinant proteins. Ponceau S (PS) stain shows protein loading. Numbers on the left indicate molecular weight marker (kDa).

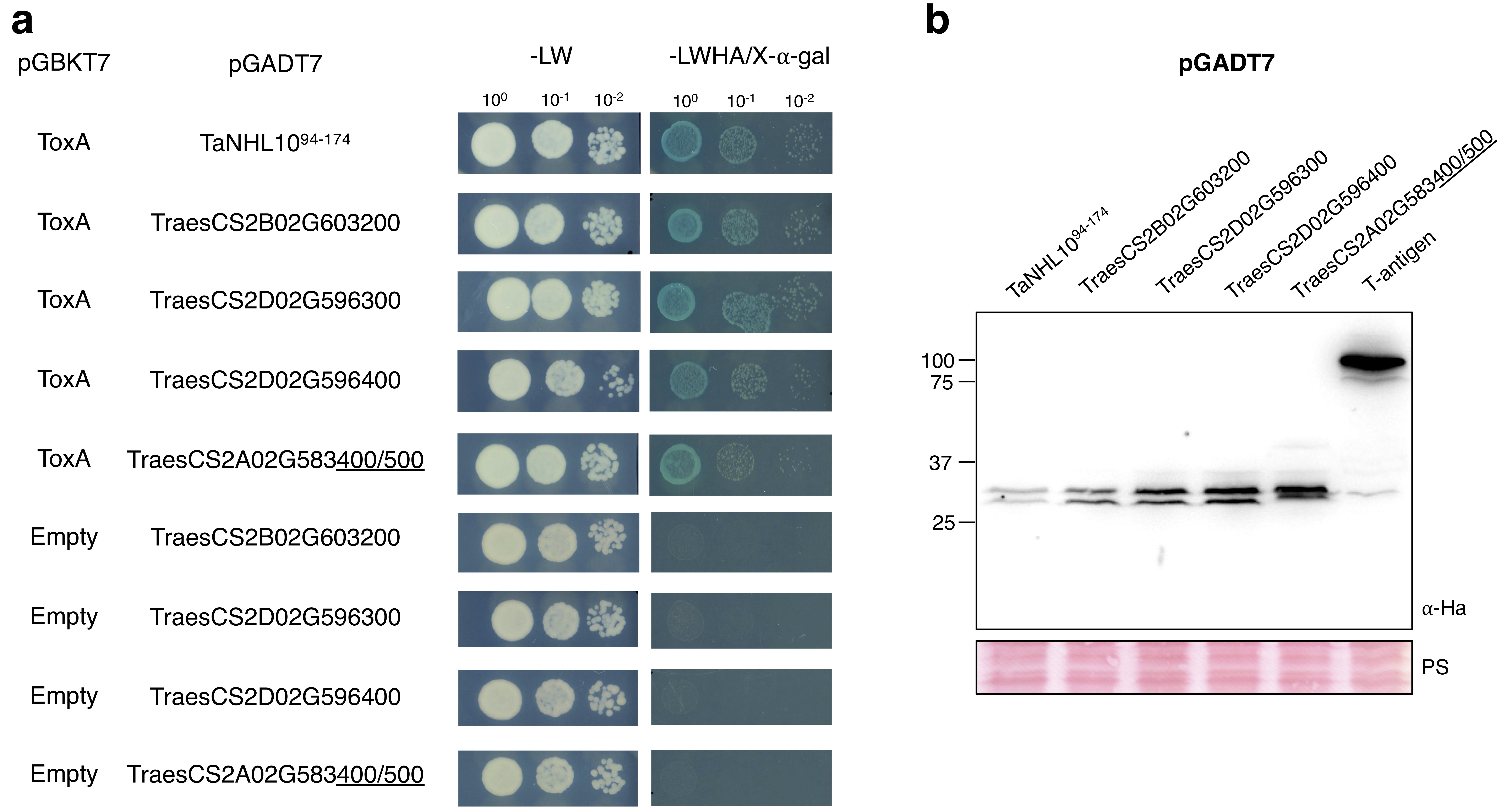

**Figure S3. Y2H assay screening ToxA interaction with five TaNHL10 homologs found in wheat genome.** Accession numbers of TaNHL10 homologs: TraesCS2B02G603200, TraesCS2D02G596300, TraesCS2D02G596400, TraesCS2A02G583400, and TraesCS2A02G583500. (a) Yeast co-expressing bait and prey plasmids were serially diluted from cell suspensions of a single yeast colony. Growth on -Leu,Trp (-LW) indicates the presence of both plasmids in the yeast colony. Growth and blue coloration on -Leu,Trp,His,Ade/X-⍺-Gal (-LWHA/X-⍺-Gal) show the interaction of two proteins. Serial dilutions reflected the strength of the interaction. ToxA was co-transformed with partial TaNHL10 (TaNHL10^94-174^), or partial TaNHL10 homologs (94-174aa). (b) Confirmation of TaNHL10 homologs (27.4 kDa) protein expression in yeast. The proteins were detected with ⍺-ha. Asterisks indicate the expected size of the recombinant proteins. Ponceau S (PS) stain shows protein loading. Numbers on the left indicate molecular weight marker.

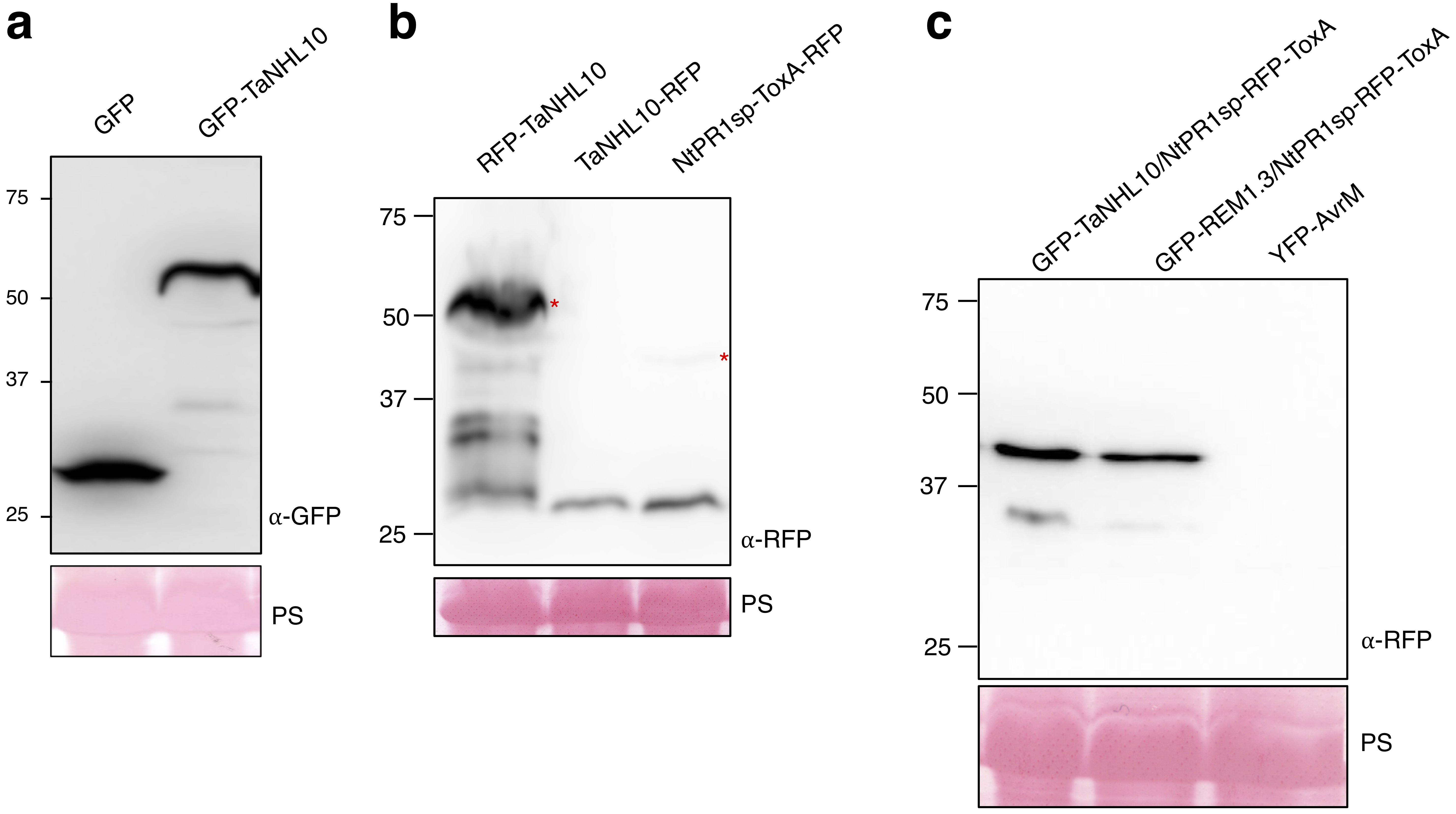

**Figure S4. Confirmation of protein expression in *N. benthamiana*.** (a) GFP (26 kDa) and GFP-TaNHL10 (51 kDa) were detected with ⍺-GFP, and (b) RFP-TaNHL10 (52 kDa), TaNHL10-RFP (52 kDa), and (c) NtPR1sp-ToxA-RFP (45 kDa) were detected with ⍺-RFP. Asterisks indicate the expected size of the recombinant proteins. Ponceau S (PS) stain shows protein loading. Numbers on the left indicate molecular weight marker (kDa).

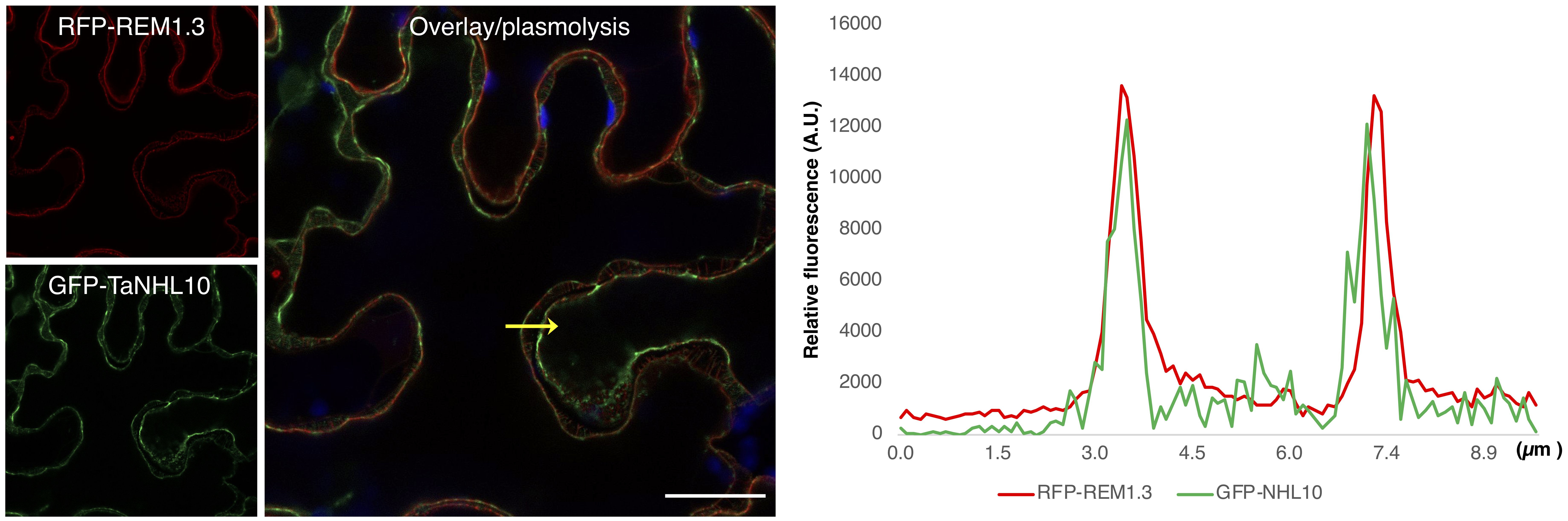

**Figure S5. TaHL10 localizes to the plasma membrane.** *N. benthamiana* plants were transiently co-expressed by agro-infiltration using the following constructs: RFP-REM1.3 (pK7WGR2/StREM1.3) and GFP-TaNHL10 (pK7FWG2/TaNHL10). The plasmolysis of the leaf cells was induced before imaging. Fluorescence intensities of RFP/GFP in membrane transects (yellow arrowheads) at 2-dpi. Chloroplast autofluorescence is in blue in overlay images. Scale bars represent 5 µm.

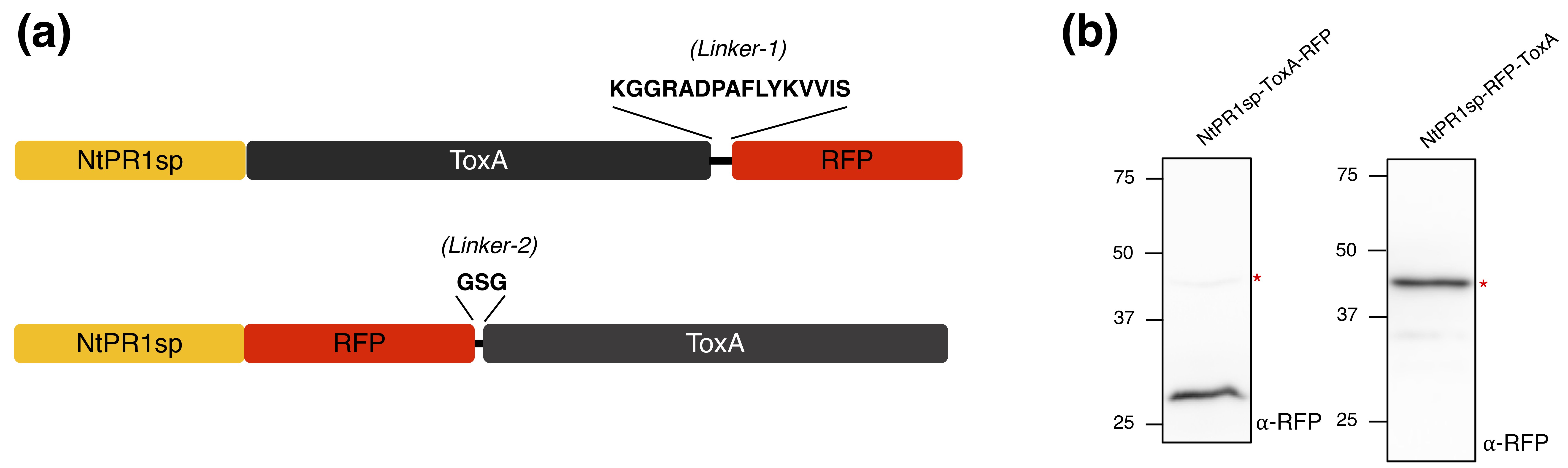

**Figure S6. Short linker prevents the cleavage of RFP tag from RFP-ToxA in apoplast.** (a) The schematic views of the secreted ToxA-RFP with linker-1 (-KGGRADPAFLYKVVIS-) and secreted RFP-ToxA with linker-2 (-GSG-). (b) Western blot analysis of expression of NtPR1sp-ToxA-RFP and NtPR1sp-RFP-ToxA constructs in *N. benthamiana* using leaf disks. Asterisks indicate the expected size of the recombinant proteins, which were detected with ⍺-RFP.

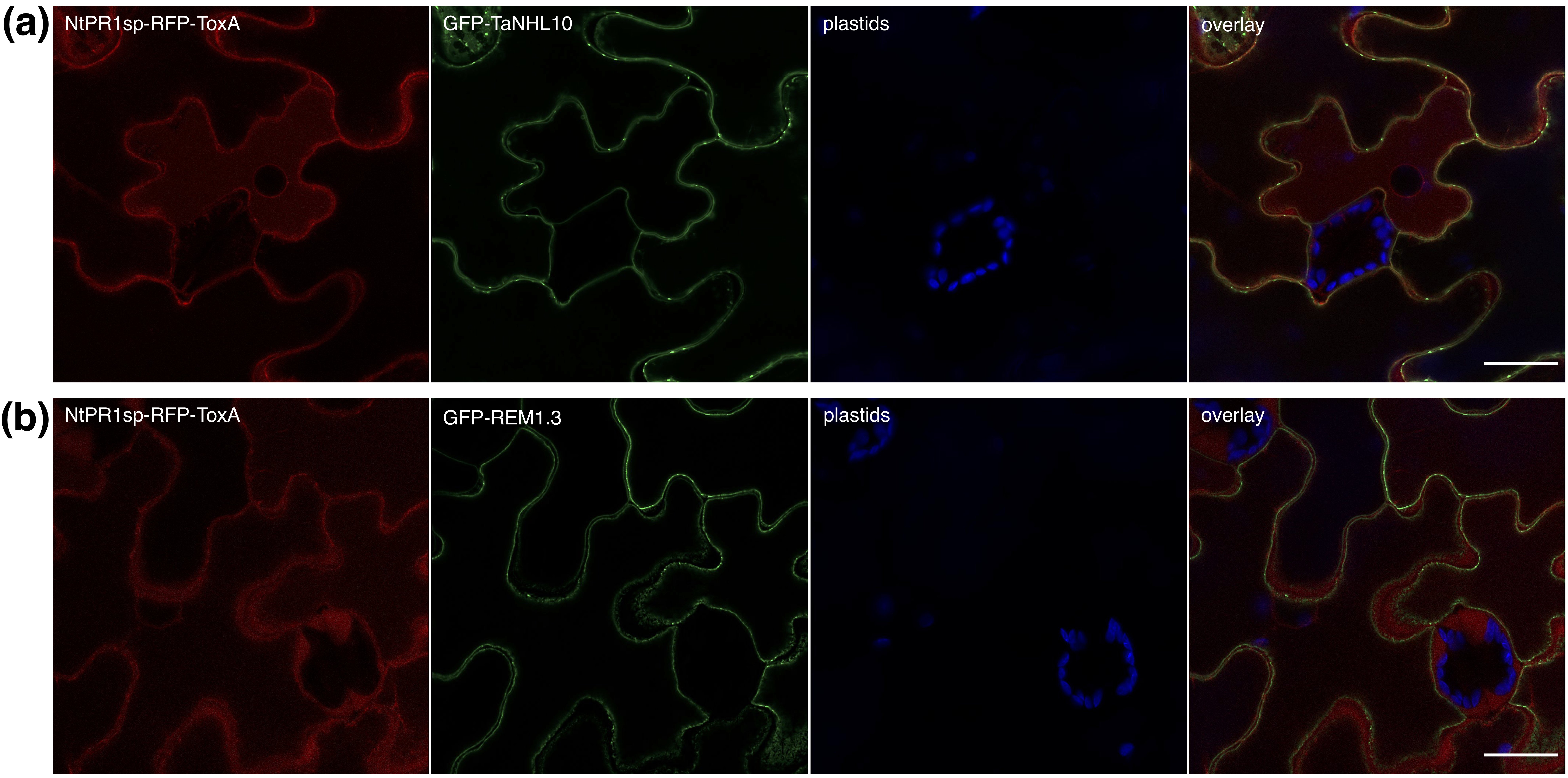

**Figure S7. Secreted-ToxA co-localizes with TaNHL10 *in* *planta*.** *N. benthamiana* plants were transiently co-expressed by agro-infiltration using the following constructs: (a) GFP-TaNHL10 and NtPR1sp-RFP-ToxA (pK7FWR2/NtPR1-mRFP-ToxA); and (b) GFP-REM1.3 and NtPR1sp-RFP-ToxA. The leaf samples were induced plasmolysis. Fluorescence intensities of RFP/GFP in membrane transects (yellow arrowheads) at 2-dpi. Chloroplast autofluorescence is shown in blue. Scale bars represent 20 µm.

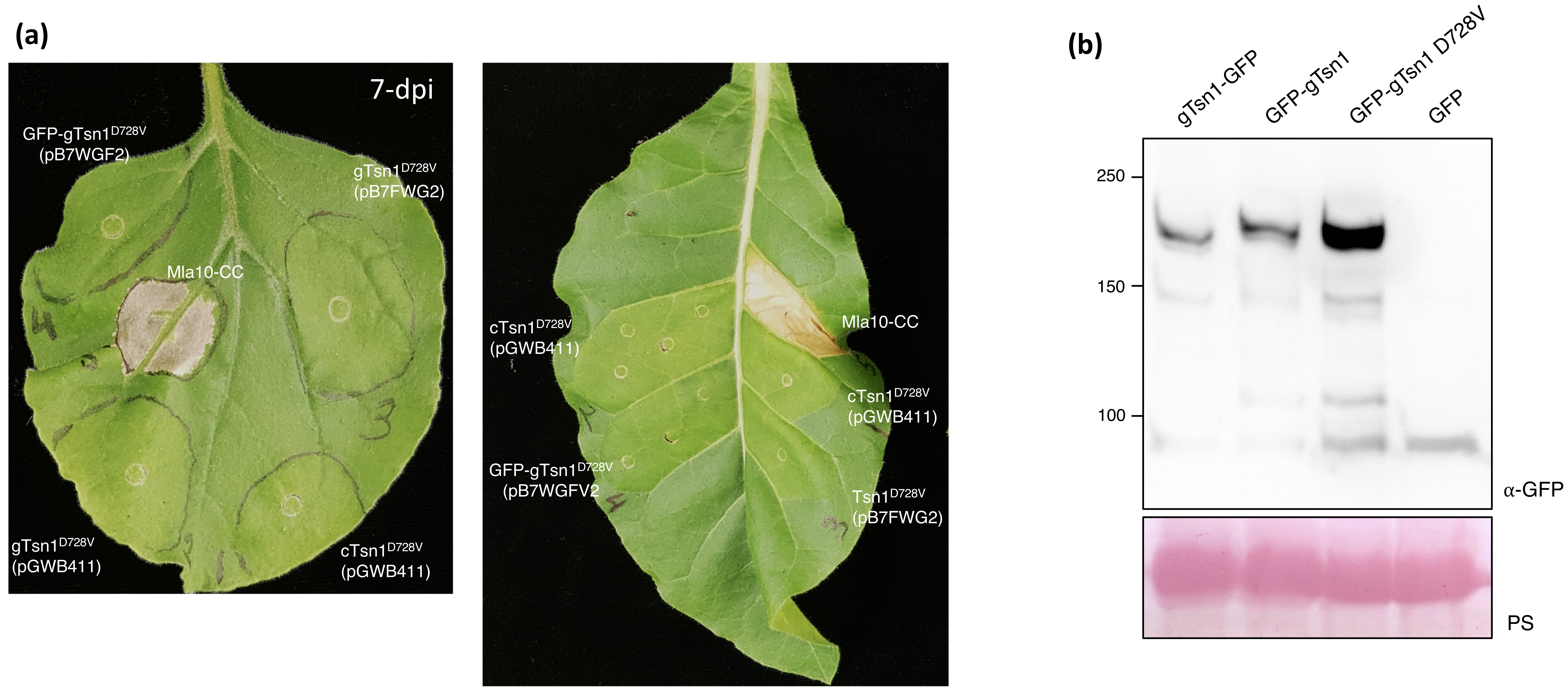

**Figure S8. The autoactive mutant of Tsn1 protein is not active in model plant systems.** (a) The genomic (gTsn1) or cDNA (cTsn1) sequence of autoactive *Tsn1* expressed in *N. benthamiana* and *N. tobaccum*. Shown are representative *N*. *benthamiana* (on the right) *and N. tobaccum* (on the left) leaf expressing autoactive Tsn1 from various constructs, 7-day post agroinfiltration. Mla CC domain expressed as positive control for cell-death development. (b) Confirmation of gTsn1-GFP, GFP-gTsn1 and GFP-gTsn1 D728V proteins expression in *N. benthamiana*. The expected size of GFP-fused Tsn1 proteins is ~190 kDa. The proteins were detected with ⍺-GFP. The GFP-expressing *N. benthamiana* samples were used for background control. Ponceau S (PS) stain shows protein loading. Numbers on the left indicate molecular weight marker (kDa).

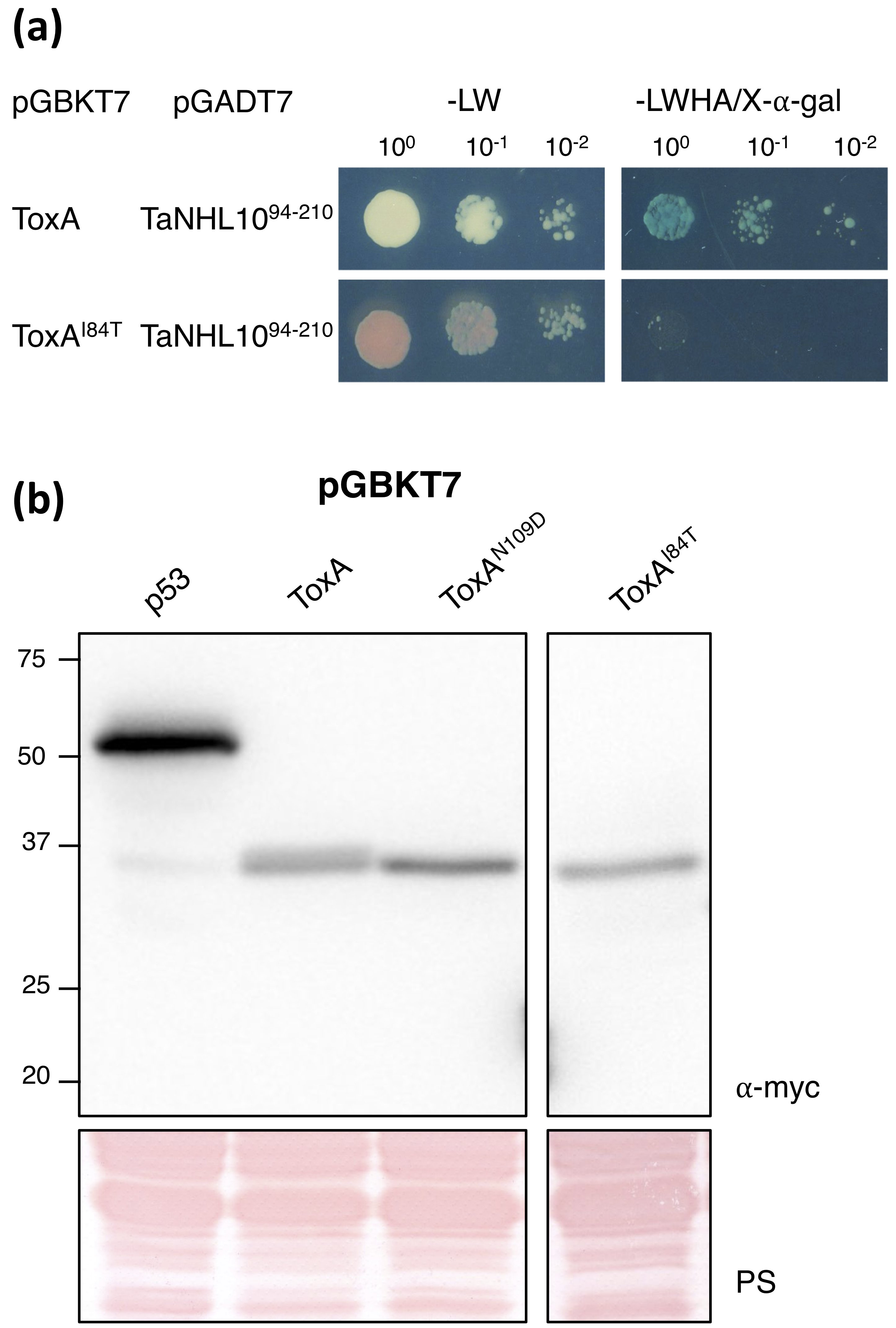

**Figure S9. A single point mutation in ToxA abolishes interaction with TaNHL10 as well as recognition by *Tsn1*.** (a) Yeast-two-hybrid assays using ToxA, ToxA^I84T^, and TaNHL10^94-210^. Yeast co-expressing bait (pGBKT7) and prey (pGADT7) plasmids were serially diluted from cell suspensions of a single yeast colony. Growth on -Leu,Trp (-LW) indicates the presence of both plasmids in the yeast colony. Growth and blue coloration on -Leu,Trp,His,Ade/X-⍺-Gal (-LWHA/X-⍺-Gal) show the interaction of two proteins. Serial dilutions reflected the strength of the interaction. (b) Confirmation of recombinant ToxA (40 kDa) and its mutants (40 kDa) protein expression in yeast. The proteins were detected with ⍺-myc. The p53 expression served as background control. Ponceau S (PS) stain shows protein loading. Numbers on the left indicate molecular weight marker (kDa).

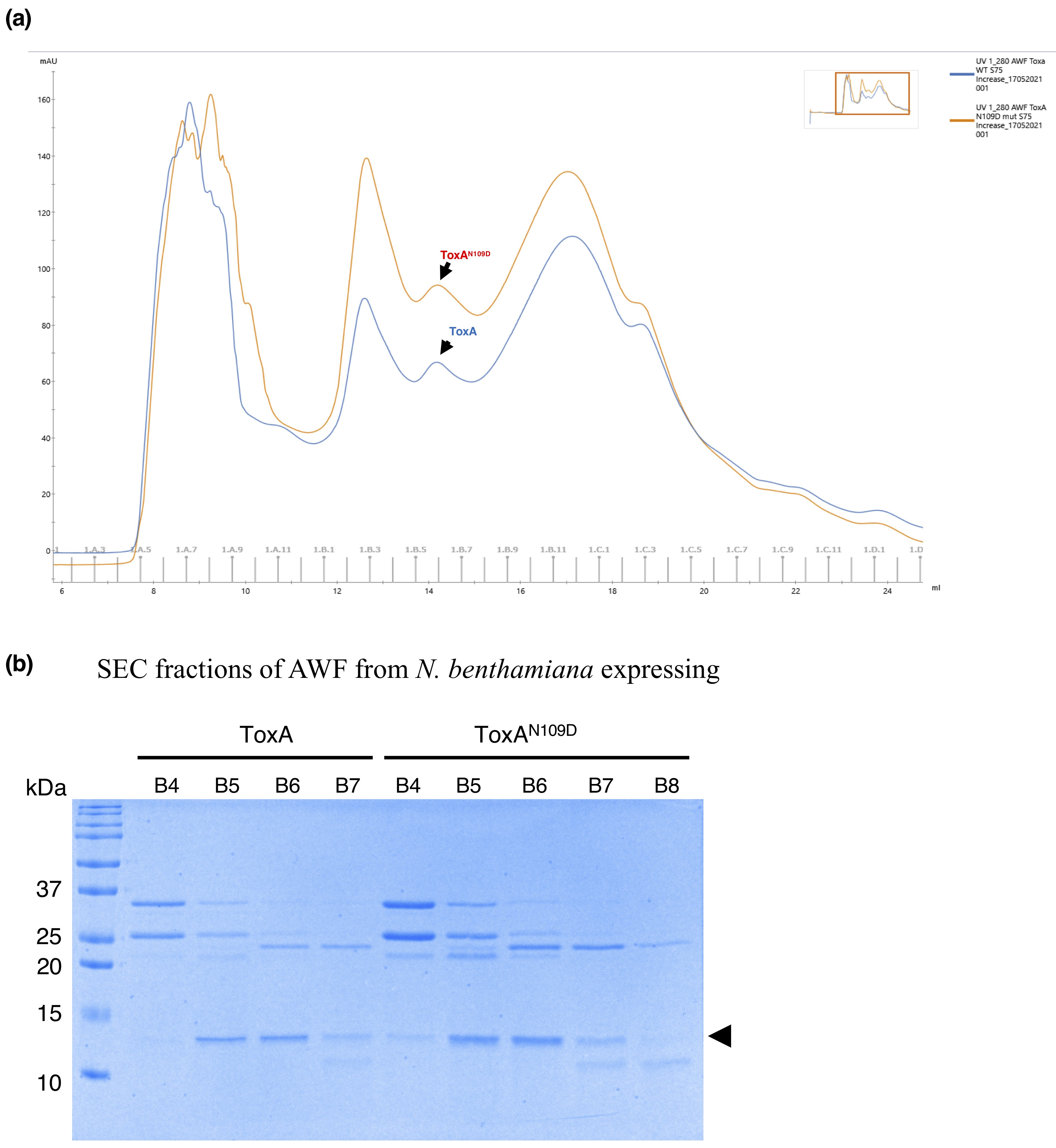

**Figure S10. Size exclusion chromatography of AWFs from *N. benthamiana* producing ToxA and ToxA^N109D^ proteins.** (a) Size exclusion chromatograms for ToxA and ToxA^N109D^ protein samples from *N. benthamiana* AWFs. (b) Coomassie-stained SDS-PAGE analysis of the SEC fractions from ToxA and ToxA^N109D^ AWF samples from *N. benthamiana,* showing ToxA^N109D^ migrated same as wild type ToxA. The black arrow indicates the expected size of the mature ToxA and ToxA^N109D^ proteins.

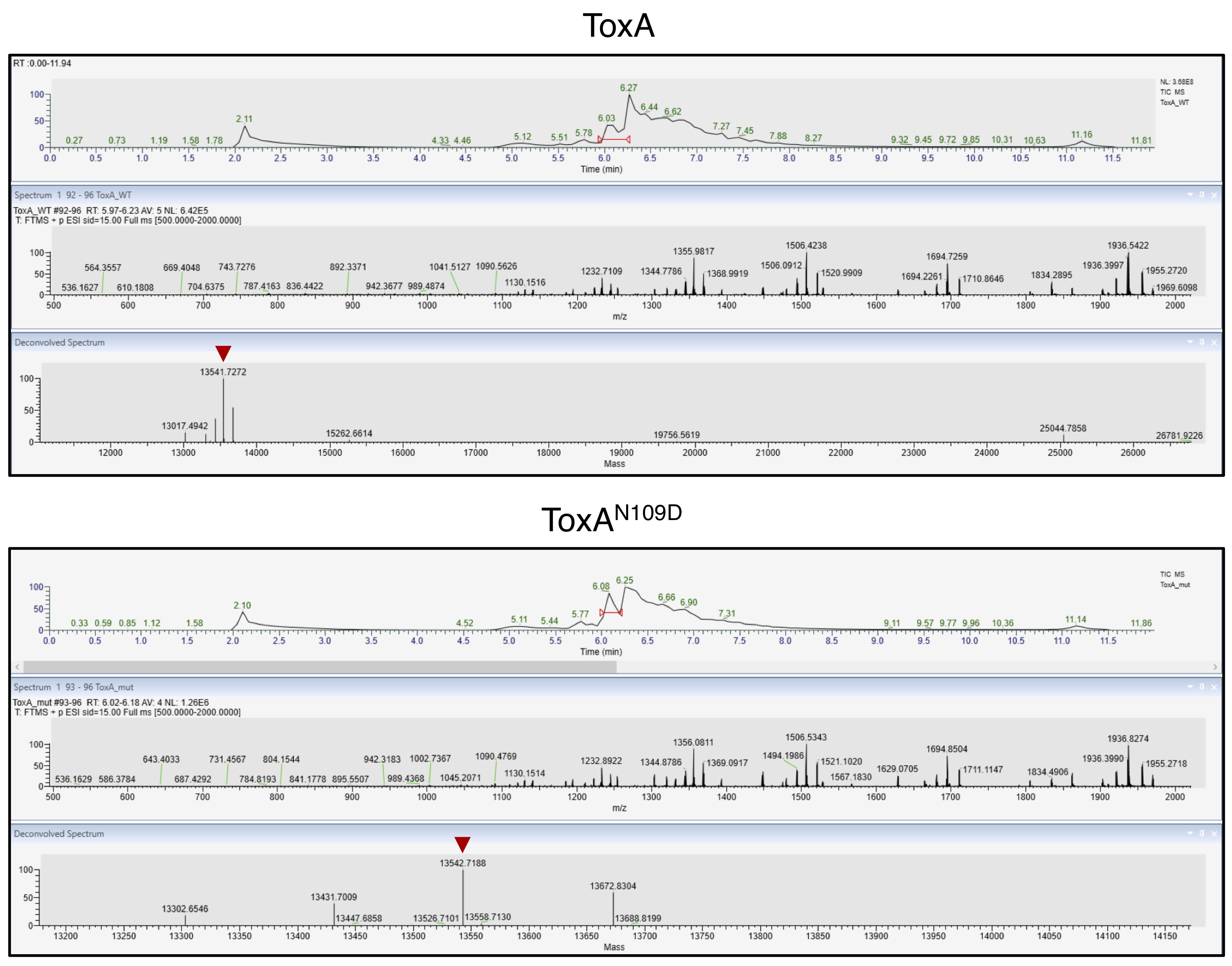

**Figure S11. Intact ToxA and ToxA^N109D^ mass spectrometry analysis.** The protein samples analysed with mass spectrometry using TripleTof mass spectrometer. For each protein, the top panel shows the elution profile from an Agilent C18 trap column; middle panel shows mass/charge ionisation series; and bottom panel shows protein reconstructed from highlighted ionisation series in the middle panel. Red arrows shows that observed masses of the proteins.

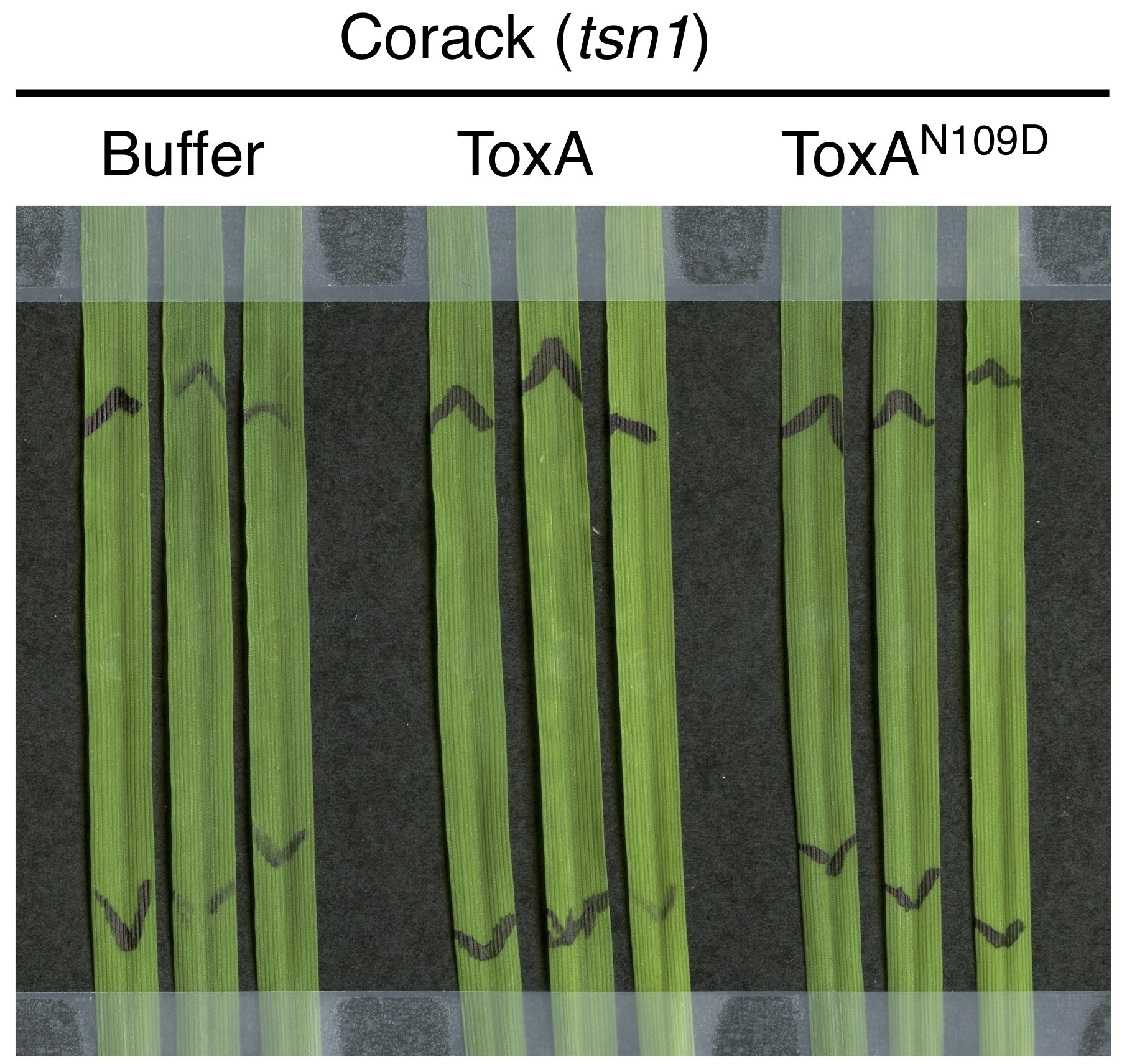

**Figure S12.** **Genotype specificity of ToxA and ToxA^N109D^ samples** (presented in Figure 5d-e). Syringe-infiltration of buffer control (10mM HEPES pH 7.5, 150 mM NaCl), ToxA, and ToxA^N109D^ samples into the second leaf of 2-week-old Corack (*tsn1*). The black markings on the leaves indicate the infiltration zone. Leaves were harvested 3-day post infiltration and imaged.

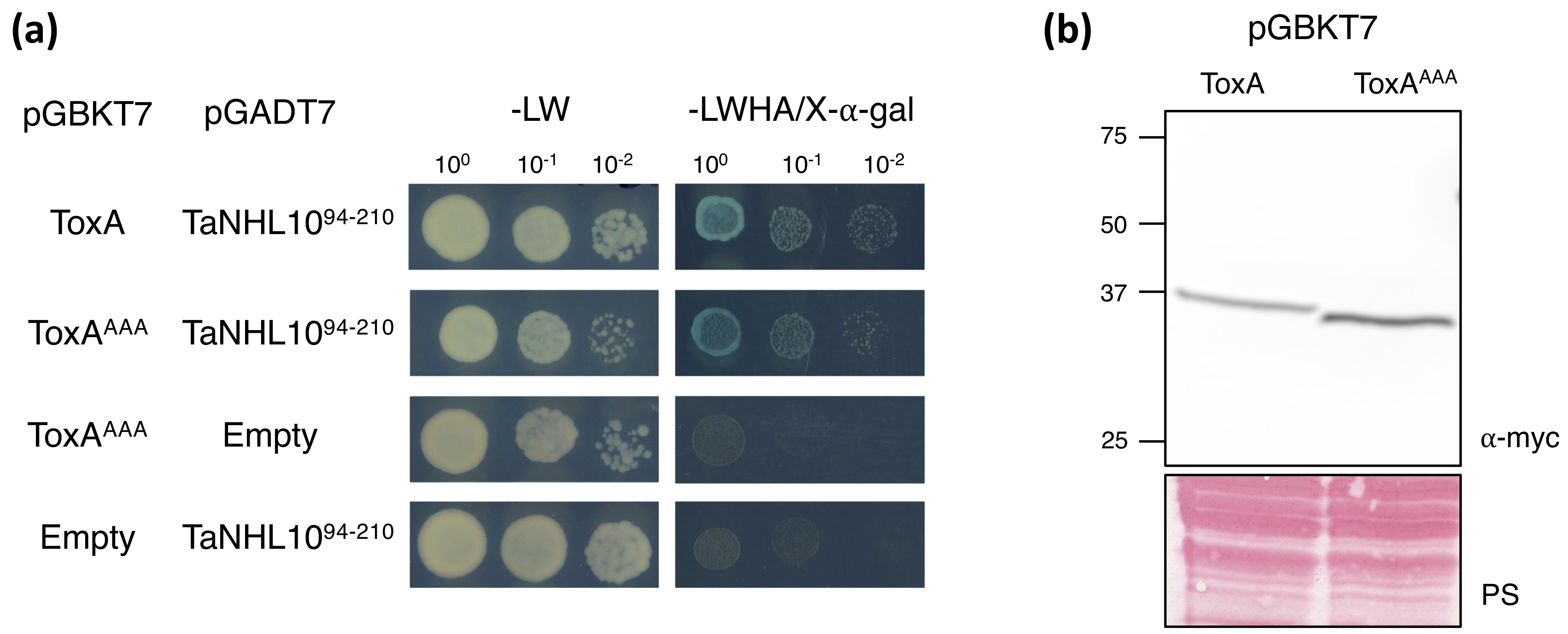

**Figure S13. The ToxA RGD motif is not required for ToxA-TaNHL10 interaction.** (a) Yeast two-hybrid assay for ToxA^AAA^ (RGD to AAA) mutant for interaction with TaNHL10^94-210^. Yeast co-expressing bait (pGBKT7) and prey (pGADT7) plasmids were serially diluted from cell suspensions of a single yeast colony. Growth on -Leu,Trp (-LW) indicates the presence of both plasmids in the yeast colony. Growth and blue coloration on -Leu,Trp,His,Ade/X-⍺-Gal (-LWHA/X-⍺-Gal) show the interaction of two proteins. Serial dilutions reflected the strength of the interaction. TaNHL10^94-210^ served as a positive control. ToxA^AAA^ and TaNHL10^94-210^ were also co-transformed with empty vector to test their autoactivity. (b) Confirmation of ToxA (40 kDa) and ToxA^AAA^ (40 kDa) protein expression in yeast. The proteins were detected with ⍺-myc. Ponceau S (PS) stain shows protein loading. Numbers on the left indicate molecular weight marker (kDa).

**Supplementary tables**

**Table S1. Primers used in this study.**

| **Primer name** | **Sequence (5’ to 3’)** |
| --- | --- |
| ToxA-gate-F | CACCATGCGTTCTATCCTCGTACTT |
| ToxA-gate-R | CTAATTTTCTAGCTGCATTCT |
| TaNHL10-F | CACCATGGGTTCGGGGAGCCGC |
| TaNHL10-3`UTR-R | CAACGGCGCAATCGCAATCGG |
| TaNHL10-R | TCAGAACCAGACTTTGCACTCG |
| TaNHL10-ST-R | GAACCAGACTTTGCACTC |
| TaNHL10-EcoR-F | TTAAGAATTCATGGGTTCGGGGAGCCGC |
| TaNHL10-20-BamHI-R | TATAGGATCCTCAGCACGCCAGGCACTTGAA |
| TaNHL10-44-EcoRI-F | TGTGGAATTCCTATACTTCATCTTCCGCCC |
| TaNHL10-BamHI-R | TATAGGATCCTCAGAACCAGACTTTGCACTC |
| TaNHL10-94-EcoRI-F | TATAGAATTCGGGCTCTACTACGACAACG |
| TaNHL10-174-BamHI-R | TTAAGGATCCTCAGAAGGCCCACACCTTGAC |
| TaNHL10-175-EcoRI-F | TATAGAATTCAAGGTGCCCGGGCCCA |
| TaPR1-9-5UTR-F | CTCACTCACTTTTGTAAGGT |
| TaPR1-9-3UTR-R | CTGCTACGTACATGCACGC |
| TaPR1-9-SP-F | CACCATGCAGAACTCGCCGCAGGAC |
| TaPR1-9-R | CTAGTATGGGCTCTGCCCCAC |
| ToxA-N109D-mut-F | CTTTGTTACCATTGGATTGGACCGCGTAAACGCC |
| ToxA-N109D-mut-R | GGCGTTTACGCGGTCCAATCCAATGGTAACAAAG |
| ToxA-I84T-mut-F | GGCCAAGTCGACACTGACAGTGTTATACTCGG |
| ToxA-I84T-mut-R | CCGAGTATAACACTGTCAGTGTCGACTTGGCC |
| Tsn1-mut-D728V-F | GTTTGTAATGCATGTCCTAGTGCACGATCTTGC |
| Tsn1-mut-D728V-R | GCAAGATCGTGCACTAGGACATGCATTACAAAC |

**Table S2. Intact mass analysis of ToxA and ToxA^N109D^ proteins.** The data is representative of the predominant species observed by intact protein-MS. The theoretical molecular weights of the proteins were predicted based on the amino acid sequences using the program PeptideMass (https://web.expasy.org/peptide_mass/), and the monoisotopic mass was used to match observable peaks in the mass spectrum. The residues of the predicted pro-domain, one amino acid following the pro-domain, and last two amino acids found at the C-terminus of the proteins are excluded in the calculation since these residues were predicted to be cleaved from the proteins in the apoplast. The observed mass of the ToxA and ToxA^N109D^ proteins should be ~2 Da less than the theoretical mass when their two cysteines are reduced.

| Proteins | Theoretical molecular weight (Da) – cysteines reduced | Theoretical molecular weight (Da) – cysteines in disulfide | Observed molecular weight (Da) |
| --- | --- | --- | --- |
| ToxA (56-170 aa) | 13543.90 | 13541.90 | 13541.73 |
| ToxA^N109D^ (56-170 aa) | 13544.88 | 13542.88 | 13542.72 |
